# Spatiotemporal organization of dynein in bulk cytoplasm promotes aster growth and positioning in large embryos

**DOI:** 10.64898/2026.04.02.716240

**Authors:** Ai Kiyomitsu, Luolan Bai, Timothy J. Mitchison, Tomomi Kiyomitsu

## Abstract

During cleavage in vertebrates, large microtubule asters grow from centrosomes in anaphase. The microtubule motor dynein pulls on these asters in bulk cytoplasm and positions centrosomes to determine cleavage geometry. However, how aster growth and pulling are coordinated remains unclear. We discovered that small metaphase asters generate a halo-like enrichment of cytoplasmic dynein at the aster periphery in medaka early embryos. In anaphase, dynein relocates from the halo to growing asters and neighboring membranous organelles, coincident with aster expansion and centrosome movement. Dynein inhibition or localized halo disruption prevents centrosome movement. Unexpectedly, dynein inhibition also suppresses aster growth and causes ectopic cytoplasmic microtubule nucleation, which leads to ectopic furrows. We propose that inactive dynein that accumulated at the metaphase aster periphery is activated in anaphase and incorporates both microtubule nucleators and organelles into growing asters to coordinately promote aster growth and pulling for efficient centrosome positioning in rapidly-dividing, large embryos.

## Introduction

Vertebrate zygotes and early embryonic blastomeres divide symmetrically for clonal expansion^1–3^, like small somatic cultured cells^4^. To achieve symmetric cell division, mitotic spindles assemble in cell centers, and equally partition chromosomes, centrosomes, and cytoplasm to daughter cells^5,6^. In typical small animal cells, astral microtubules (MTs) emanating from centrosomes, reach the cell cortex and interact with cortical dynein complexes^7,8^, which pull on astral MTs to correct spindle misposition to achieve spindle centering^4,9,10^. In contrast, astral MTs do not reach the mitotic cell cortex in large embryos due to a scaling mismatch between spindle and cell size^1–3,11–14^. Thus, centrosomes are instead centered before mitotic entry in large embryos^1^. To achieve centrosome centering, each embryonic centrosome specifically grows a large MT meshwork called the “aster”^1^. Asters dramatically expand in anaphase and promote centrosome movement toward the center of daughter blastomeres throughout cytokinesis^1,3^. Previous research indicates that the centrosomes are centered by length-dependent aster-pulling by cytoplasmic dynein in large frog and fish embryos^1,2,15^.

Cytoplasmic dynein-1 (hereafter referred to as dynein) is a conserved, minus-end-directed, MT-based motor that has multiple, essential functions^16^, including organelle transport^17^, spindle assembly^18–21^, and MT-dependent force generation^1,10,15,22,23^. Recent structural studies have established that dynein is auto-inhibited^24^, but it is activated by interacting with dynactin and activating adaptors^25,26^ that connect various cargos with dynein-dynactin for dynein-driven transport and force-generation. In addition to aster pulling in bulk cytoplasm^1^, dynein functions in pronucleus positioning in mouse zygotes^27^, parental genome clustering in human and bovine zygotes^28^, and centrosome-pronucleus interaction and centrosome pulling in *C. elegans* one-cell embryos^22,29^. However, it remains unclear how the localization and activity of dynein are spatiotemporally regulated, especially during the aster-pulling process in large vertebrate cytoplasm.

Asters consist of many short, interacting MTs^30^. Recent studies with egg extract and mathematical modeling propose a two-step model for large aster growth^31^. First, centrosome-dependent MT nucleation initiates aster growth in early mitosis, which forms stationary asters having short radii in metaphase. Subsequently, in response to anaphase onset, asters dramatically increase their radii via autocatalytic MT-stimulated MT nucleation away from centrosomes. This model explains aster growth in frog and fish embryos^1^. However, how dynein contributes to aster growth remains poorly investigated in large embryos.

In this study, we visualized endogenous dynein and dynactin using CRISPR in live medaka fish embryos and analyzed their functions using chemical, genetic, and mechanical perturbations to understand their contribution to aster growth and positioning. Unexpectedly, we found that dynein and dynactin show a halo-like cytoplasmic enrichment around the metaphase aster periphery that assists both aster expansion and positioning in daughter blastomeres.

## Results

### A dynein halo surrounds metaphase spindles in medaka embryos

We first visualized endogenous dynein in medaka early embryos. Using CRISPR/Cas9, we generated three knock-in strains expressing either dynein heavy chain (DHC)-mCherry (DHC-mCh), DHC-mClover-3xFLAG (DHC-CF), or DHC-mAID-mClover-3xFLAG (DHC-mACF) (Fig. S1A-C)^3,32^. All homozygous embryos were viable and showed identical localization of dynein. We next compared localization of dynein (DHC-mCh) with that of MTs (EGFP-α-tubulin) in early embryonic division. As observed in zebrafish embryos^1,33^, radial arrays of interphase MTs were disassembled before mitotic entry (Fig. 1A-B), and reorganized by dominant aster growth after anaphase (Fig. 1B-C). Unexpectedly, dynein displayed novel, cell-cycle-dependent localization dynamics that differed from those of MTs and previous reports of cortex-associated dynein^4,9,10,34,35^. In interphase, dynein accumulated around centrosomes and nuclei (Fig. 1B). However, upon mitotic entry, dynein was reduced around centrosomes and in cytoplasm near asters (Fig. 1B, Fig. S2A-C). Importantly, dynein showed a halo-like cytoplasmic enrichment around the periphery of metaphase asters (Fig. 1B arrows, Fig. 1D-F), which we call the dynein “halo” based on its appearance (Movie S1 and S2). To carefully analyze the spatial relationship between astral MTs and the dynein halo, we next visualized MTs with the MT-binding domain of ensconsin fused to three GFPs (EMTB-3xGFP) (Fig. 1G-H, Fig. S3A-B)^1,36^. Aster edge was located around the inner surface of the dynein halo in metaphase (Fig. 1G-H). After anaphase onset, the dynein halo moved slightly outward, coupled with aster expansion (Fig. 1C, 1H t=3). In parallel, dynein appeared to relocate from the halo to asters and to move toward centrosomes (Fig. 1F, Fig. 1H), showing the original centrosome accumulation in interphase. Interestingly, a small population of MTs appeared to be nucleated in cytoplasm between the aster edge and the cell cortex during anaphase (Fig. 1H t=2-3, Movie S3, Fig. S3A-C), but these cytoplasmic MTs were eventually incorporated by expanding asters (Fig. 1H). The dynein halo was repeatedly observed in the subsequent cleavage cycle (Fig. 1B, Fig. S2D), but reached the cell cortex after the 8-cell stage as cells became smaller (Fig.S2D). However, dynein generally appeared to be excluded near metaphase spindles, even in small blastula-stage blastomeres (Fig. S2D). These data are consistent with the localization of dynein in sea urchin embryos^37^.

**Figure 1.**
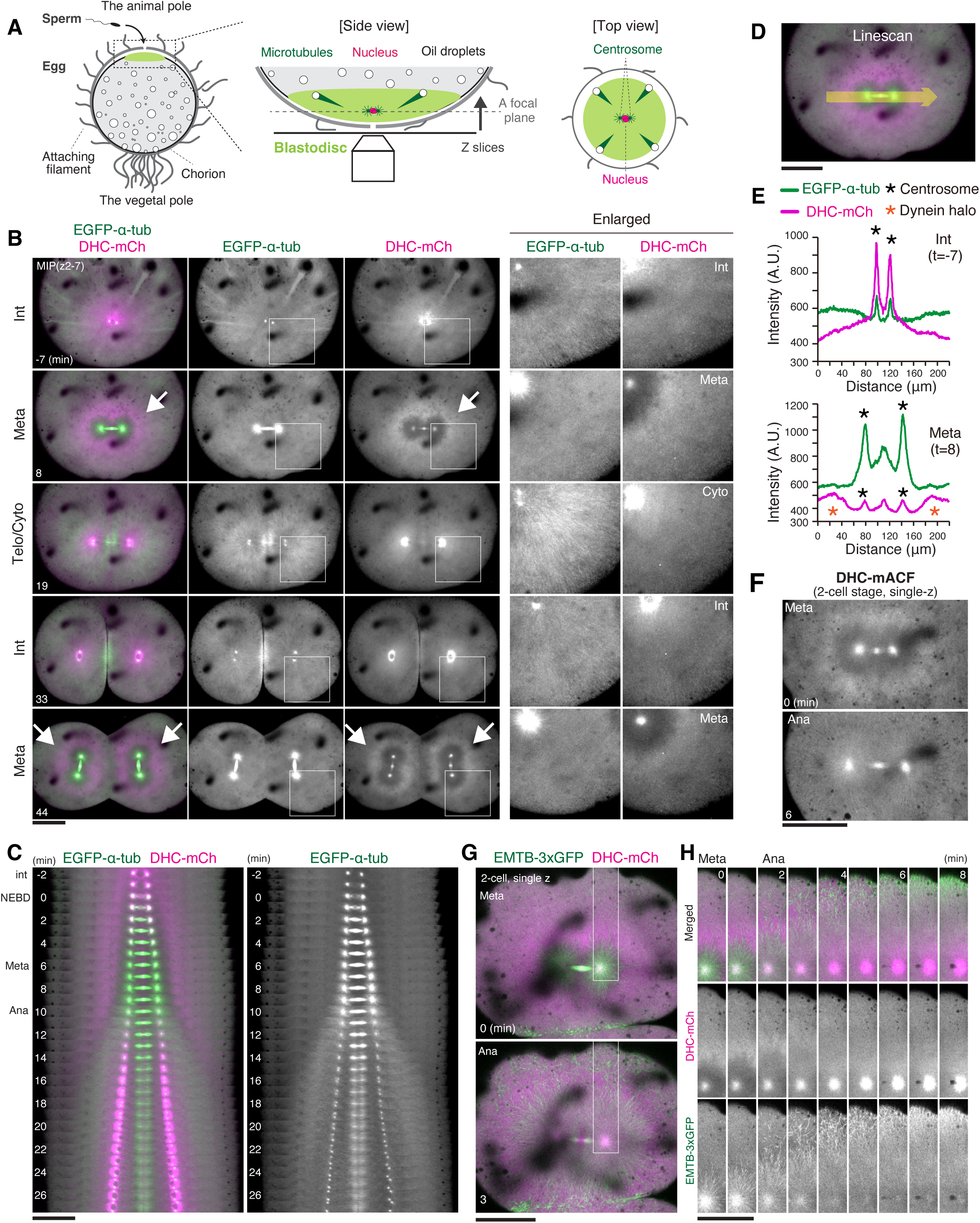
The dynein halo is formed in a cell-cycle-dependent manner during early embryonic divisions. (A) Schematic representation of microscopic observation of a medaka fertilized egg. (B) Maximum intensity projection (MIP) images of a live embryo showing cell-cycle-dependent localization dynamics of MTs (green) and dynein (magenta) during zygotic division. Right panels show enlarged images. (C) Kymographs based upon Movie S1 showing MTs (green) and dynein (magenta) during the first mitosis. (D) Fluorescence intensity on the yellow arrow (Mean, 24px width) was measured. (E) Graphs of fluorescence intensities for line scans of interphase and metaphase embryos. (F) Live embryo images showing localization changes of endogenous dynein, DHC-mACF, between metaphase and anaphase. (G) Live images showing localization of dynein (DHC-mCh) and asters by EMTB-3xGFP in metaphase and anaphase. (H) Sequential live-cell images of EMTB-3xGFP and dynein showing dissolution of the dynein halo in response to anaphase aster expansion. Scale bars = 100 μm.

As with dynein, endogenous p50 (p50-mACF) (Fig. S1D), a dynactin subunit, also displayed a halo-like enrichment in metaphase cytoplasm (Fig. S 4A-B) and cell-cycle-dependent localization change during early embryonic division (Fig. S 4A-C). Remarkably, p50 was more clearly detectable on asters after anaphase (Fig. S4B t=12), at cleavage planes, at membranes of karyomeres^38^, and at nuclear envelopes after karyomere fusion (Fig. S4B-C), probably as dynactin contains four copies of p50^39,40^. These results indicate that dynein-dynactin complexes localize at multiple subcellular sites, including the cytoplasmic halo-like zone in metaphase, and that they dynamically change their localization in a cell-cycle-dependent manner during the cleavage cycle.

### The dynein halo is created by metaphase asters

Since the dynein halo forms around the aster periphery (Fig. 1G-H), we next analyzed roles of MTs in dynein halo formation. We first visualized chromosomes (RCC1-mCh) and dynein (DHC-CF) in the presence of the MT-destabilization drug, nocodazole (Fig. 2A-D), which disrupts spindle formation, but does not induce mitotic delay or arrest in medaka early embryos^3^. Dynein was detected at centrosomes and the cytoplasmic halo during prometaphase and metaphase in control blastomeres (Fig. 2A), but not in those treated with nocodazole (Fig. 2C t=3-6). In controls, dynein likely localized at kinetochores during prometaphase (Fig. 2B). It also appeared to localize at spindle MTs near chromosomes in metaphase and at central spindles during anaphase (Fig. 2B), as these signals were disrupted by nocodazole-treatment (Fig. 2D). Intriguingly, dynein displayed membrane-like localization specifically in interphase around the nucleus and in cytoplasm of nocodazole-treated embryos (Fig. 2C t=0 and 21), suggesting cell-cycle dependent association of dynein with membranous organelles.

**Figure 2.**
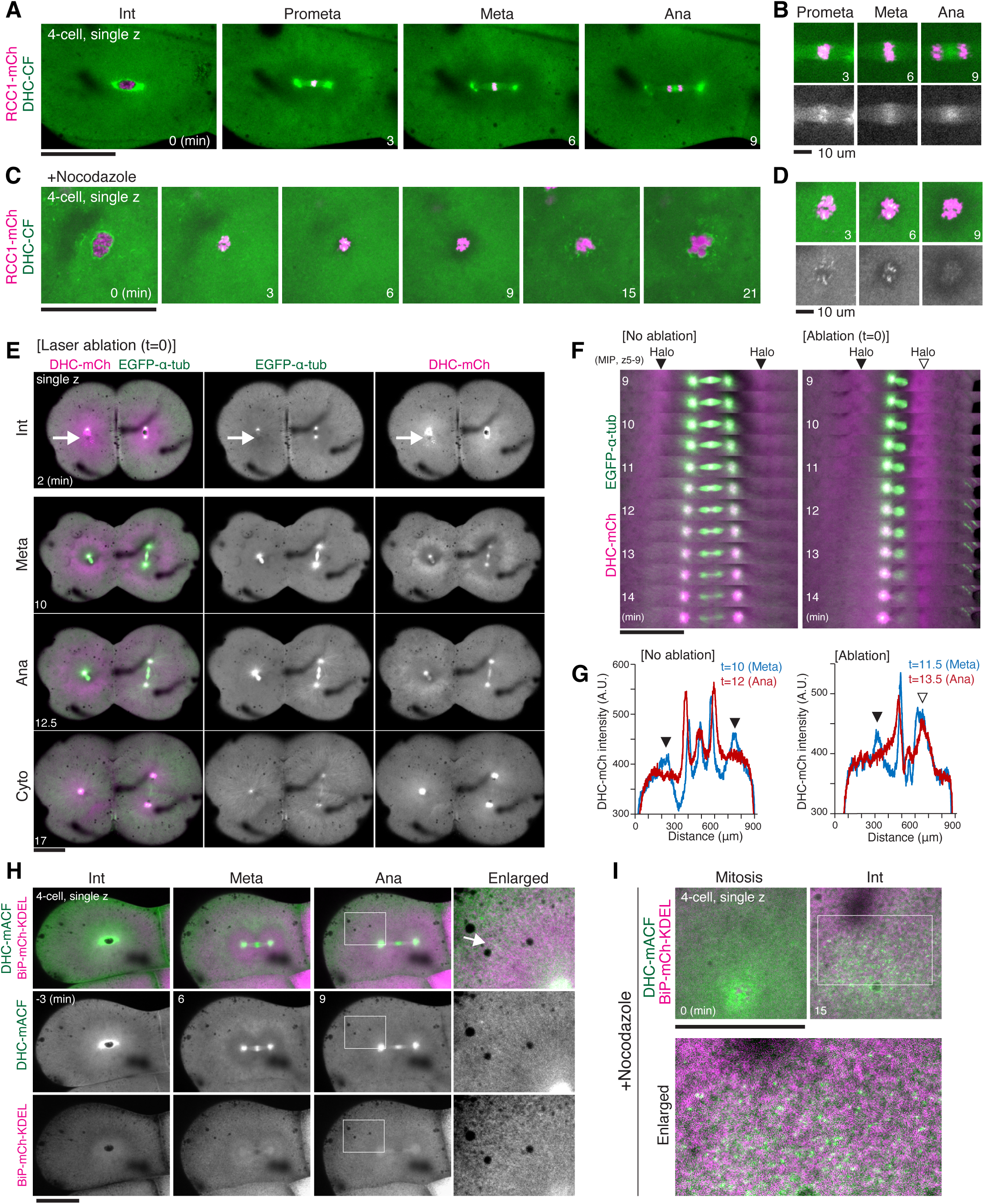
Asters are required for both formation and dissolution of the dynein halo during mitosis. (A-D) Live images showing chromosomes (RCC1-mCh) and dynein (DHC-CF) in control (A-B) and 170 nM nocodazole-treated embryos (C-D). (E) Live 2-cell-embryo images showing MTs (green) and dynein (magenta) after laser ablation. The centrosome indicated by white arrows was disrupted. (F) Kymographs showing dynein halo (magenta) dynamics in control (left) and a centrosome-ablated blastomere (right). Centrosome ablation changes the behavior of the dynein halo (a white triangle). (G) Graphs of fluorescence intensities for line scans of metaphase and anaphase embryos in control (left) and a centrosome-ablated blastomere (right). (H) Live images of dynein (DHC-mACF) and an ER marker (BiP-mCh-KDEL) showing partial co-localization in interphase and anaphase. Right: enlarged images in anaphase. An arrow indicates punctate dynein signals, likely co-localizing with ER on anaphase asters. ER tends to enrich in the dynein-less cytoplasmic region near the spindle in metaphase. (I) Live images of dynein (DHC-mACF) and an ER marker (BiP-mCh-KDEL) in 4-cell blastomeres after 2μM nocodazole-treatment. BiP-mCh-KDELs were exogenously expressed by mRNA injection in 1-cell embryos and thus their intensities increased during embryogenesis. Scale bars = 100 μm except for (B) and (D) in which scale bars = 10 μm.

To further validate the requirement of asters for dynein halo formation, we next disrupted one centrosome in a pair using laser ablation prior to 2-cell mitosis (Fig. 2E). Centrosome ablation created a mono-polar spindle with a single aster (Fig. 2E-F), which excluded dynein from the aster and enriched dynein near the aster periphery, whereas dynein was concentrated near the monopolar spindle at the opposite side in metaphase (Fig. 2E-F t=10), where it tended to remain even after anaphase (Fig. 2E-G, Fig. S5A, Movie S4). These results demonstrate that asters are required for both formation and dissolution of the dynein halo throughout mitosis.

### Dynein partially colocalizes with ER in asters during anaphase

After nocodazole treatment, dynein showed fibrous membrane-like localization specifically in interphase (Fig. 2C). To understand how dynein interacts with membranous organelles during the cell cycle, we next compared localization of dynein (DHC-mACF) with that of ER, as ER is transported and organized by dynein^17^. Both dynein and ER, visualized by ER marker (BiP-mCh-KDEL)^41^, accumulated near the nucleus in interphase (Fig. 2H), but ER tended to enrich in dynein-less cytoplasm near the spindle in metaphase (Fig. 2H, Fig. S5B-C). During anaphase, ER appeared to partially colocalize with dynein in anaphase asters (Fig. 2H, Fig. S5B-C). Nocodazole treatment produced punctate dynein signals after mitotic exit (Fig. 2I),and these dynein signals also partially colocalized with ER. These results suggest that dynein interacts with membranous organelles throughout interphase, including ER and the nuclear envelope after anaphase onset, but that it dissociates from them after mitotic entry. These data are consistent with observation in Xenopus egg extract^42^.

### Dynein inhibition causes ectopic cytoplasmic MT nucleation during anaphase in zygotes

To probe dynein’s functions, we next inhibited dynein-dynactin interaction by injecting a dominant-negative fragment of the dynactin p150 coiled-coil 1 (p150-CC1)^1,43^ just prior to mitosis in zygotes. Although 28% of injected embryos (n=7/25) did not assemble spindles (Fig. S7A-E), 72% of injected embryos (n=18/25) entered mitosis and formed zygotic spindles (Fig. 3A-B). In these p150-CC1 injected zygotes, dynein was hardly detectable at centrosomes, spindles, and halos, in contrast to control embryos (Fig. 3A-B), showing that p150-CC1 inhibits dynein localization. MT (EGFP-α-tubulin) intensities were equivalently high at asters and metaphase spindles in controls (Fig. 3A, 3C), but were greatly reduced at asters in p150-CC1 injected blastomeres (Fig. 3B, 3D).

**Figure 3.**
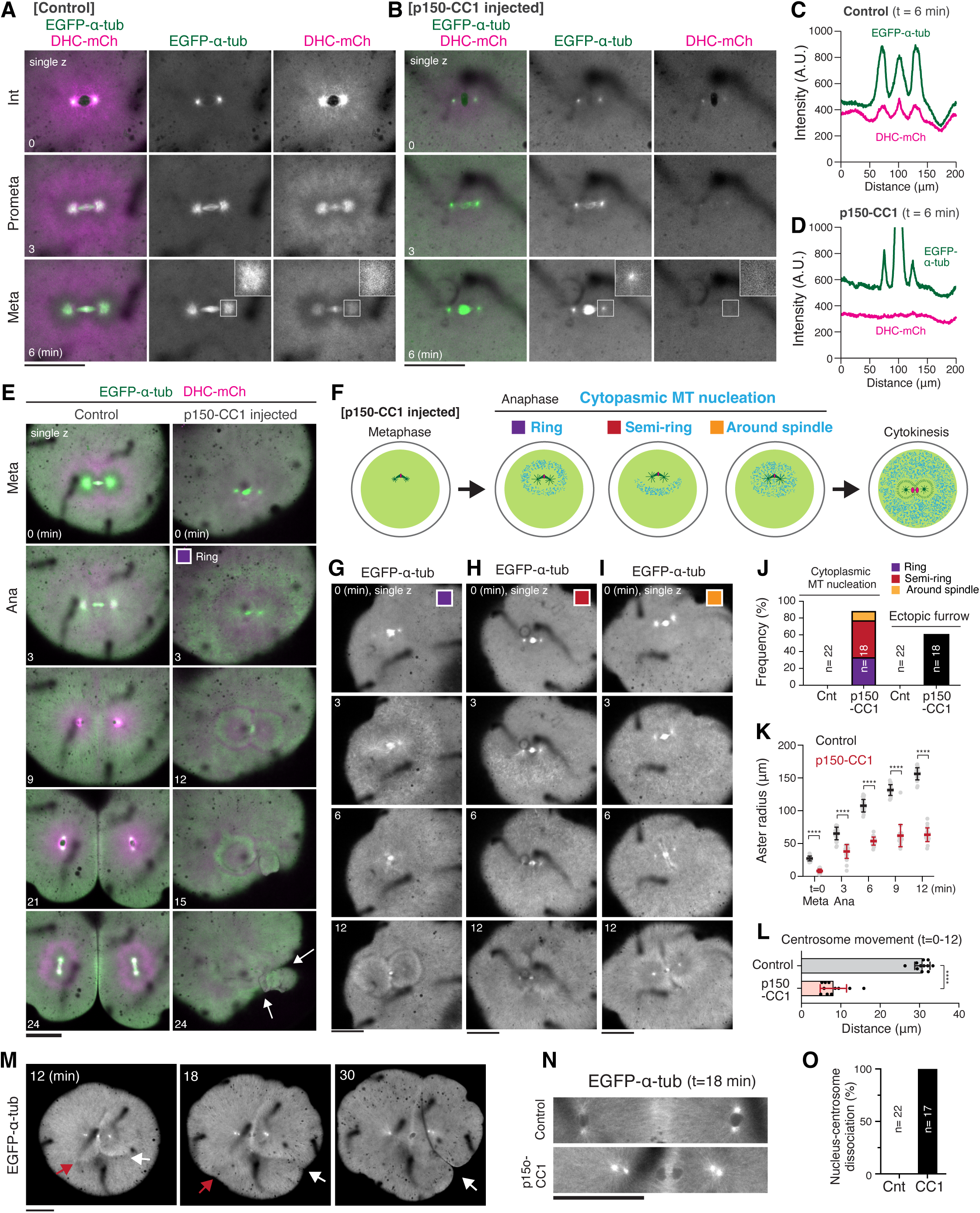
Dynein inhibition causes cytoplasmic MT nucleation in anaphase, leading to ectopic furrow formation. (A-B) Live images showing MTs (EGFP-α-tubulin, green) and dynein (DHC-mCh, magenta) in control (A) and p150-CC1 injected zygotes (B). (C-D) Graphs of fluorescence intensities for line scans (24px width) of EGFP-α-tubulin and DHC-mCh in control (C) and p150-CC1 injected (D) metaphase embryos. (E) Live images of MTs and dynein in control (left) and p150-CC1 injected zygotes (right) showing anaphase cytoplasmic MT nucleation and ectopic furrow formation (arrows) caused by p150-CC1 injection. (F) Schematics of initial patterns of cytoplasmic MT nucleation in anaphase: ring, semi-ring and around the spindle. (G-I) Representative images of cytoplasmic MT nucleation showing ring (G), semi-ring (H), or around the spindle (I) in anaphase. Intensities of EGFP-α-tubulin were adjusted differently in G-I. (J) Graphs showing frequencies of cytoplasmic MT nucleation or ectopic furrows in control (n=22) and p150-CC1 injected zygotes (n=18). (K) Graphs showing aster radii in control (n=22) and p150-CC1 injected (n=22) zygotes. ****p<0.0001 (L) Graphs showing distances of centrosome movement in control (n=11) and p150-CC1 injected zygotes (n=11). ****p<0.0001 (M) Live images of MTs showing ectopic furrow formation (white arrows) and integration of asters with cytoplasmic MTs (red arrows). See Fig. S6B and S6D for details. (N) Live images of MTs showing positions of nuclei (dark oval regions) and centrosomes during cytokinesis in control (top) and p150-CC1 injected zygotes (bottom). (O) Frequencies of nucleus-centrosome dissociation in control and p150-CC1 injected zygotes. Scale bars = 100 μm.

Unexpectedly, gross MTs were nucleated in anaphase cytoplasm in p150-CC1 injected zygotes (Fig. 3E-F, S6A-C, Movie S5, 6), a condition rarely observed in anaphase controls (Fig. 3E, S6C). These cytoplasmic MTs displayed either a ring or semi-ring shape in >75% of p150-CC1 injected embryos (n=18, Fig. 3F-J), and quickly appeared throughout the cytoplasm (Fig. 3G-I, S6C). Cytoplasmic MTs likely compete with asters for free tubulin and form a midzone-like, interaction zone at the boundary between asters and cytoplasmic MTs (Fig. 3E t=12)^2^, resulting in suppression of aster growth (Fig. 3K). Outward centrosome movement was also impaired after anaphase (Fig. 3L). During cytokinesis, normal furrow did not form in p150-CC1 injected zygotes (Fig. 3E, 3M). Instead, ectopic furrows frequently formed between boundaries of asters and cytoplasmic MTs (Fig. 3E, 3J, 3M white arrows). In some cases, asters eventually merged with cytoplasmic MTs (Fig. 3M, Fig. S8D). Other events such as nuclear migration and nucleus-centrosome interactions were also impaired in dynein inhibited embryos (Fig. 3N-O). Abnormal cytoplasmic MT nucleation was also observed in p150-CC1 injected embryos that did not form spindles (n=7/25) (Fig. S7A-F). In these embryos, cytoplasmic MTs were nucleated before nuclear envelope breakdown (NEBD) and spread from the cell periphery to cell center, resulting in abnormal furrow formation (Fig. S7B, S7D). Intriguingly, NEBD failed to occur in 4 of 7 embryos (Fig. S7E), consistent with the role of dynein in NEBD^44,45^. Together, these results indicate that dynein is required to integrate cytoplasmic MTs or MT nucleators into anaphase asters and/or suppress ectopic cytoplasmic MT nucleation to promote aster expansion and proper cytokinesis in large medaka zygotes.

### Dynein halo disruption causes cytoplasmic MT nucleation and centrosome positioning defects

To validate dynein inhibition phenotypes, we next injected p150-CC1 into one blastomere of 2-cell embryos (Fig. 4A). Consistent with results in zygotes, p150-CC1 injection disrupted the dynein halo and dynein’s centrosome localization at metaphase (Fig. 4A-D, Fig. S8A), leading to ectopic cytoplasmic MT nucleation in anaphase (Fig. 4B, S8B). Strikingly, these cytoplasmic MTs formed Y-shaped MT-less regions between cytoplasmic MTs and asters (Fig. 4B t=9, Fig. S8B) and hindered aster growth near cytoplasmic MTs (Fig. S8C). Importantly, most asters subsequently elongated and formed bud-like ectopic furrows (Fig. 4B, Fig. S8B, Movie S7), suggesting that asters can elongate in the absence of functional dynein. However, these growing asters were insufficient to form cleavage furrows between asters (Fig. 4B t=18, Fig. 4D, Fig. S8D) or to move centrosomes outward in p150-CC1 injected blastomeres (Fig. S8E). We found that centrosome movement was also attenuated in adjacent non-injected blastomeres (Fig. S8E), most likely due to leakage of p150-CC1 to sister blastomeres. Consistently, non-injected sister blastomeres showed abnormal centrosome positioning before onset of 4-cell mitosis (Fig. S8F), resulting in abnormal spindle formation (Fig. 4B t=24, Fig. S8F).

**Figure 4.**
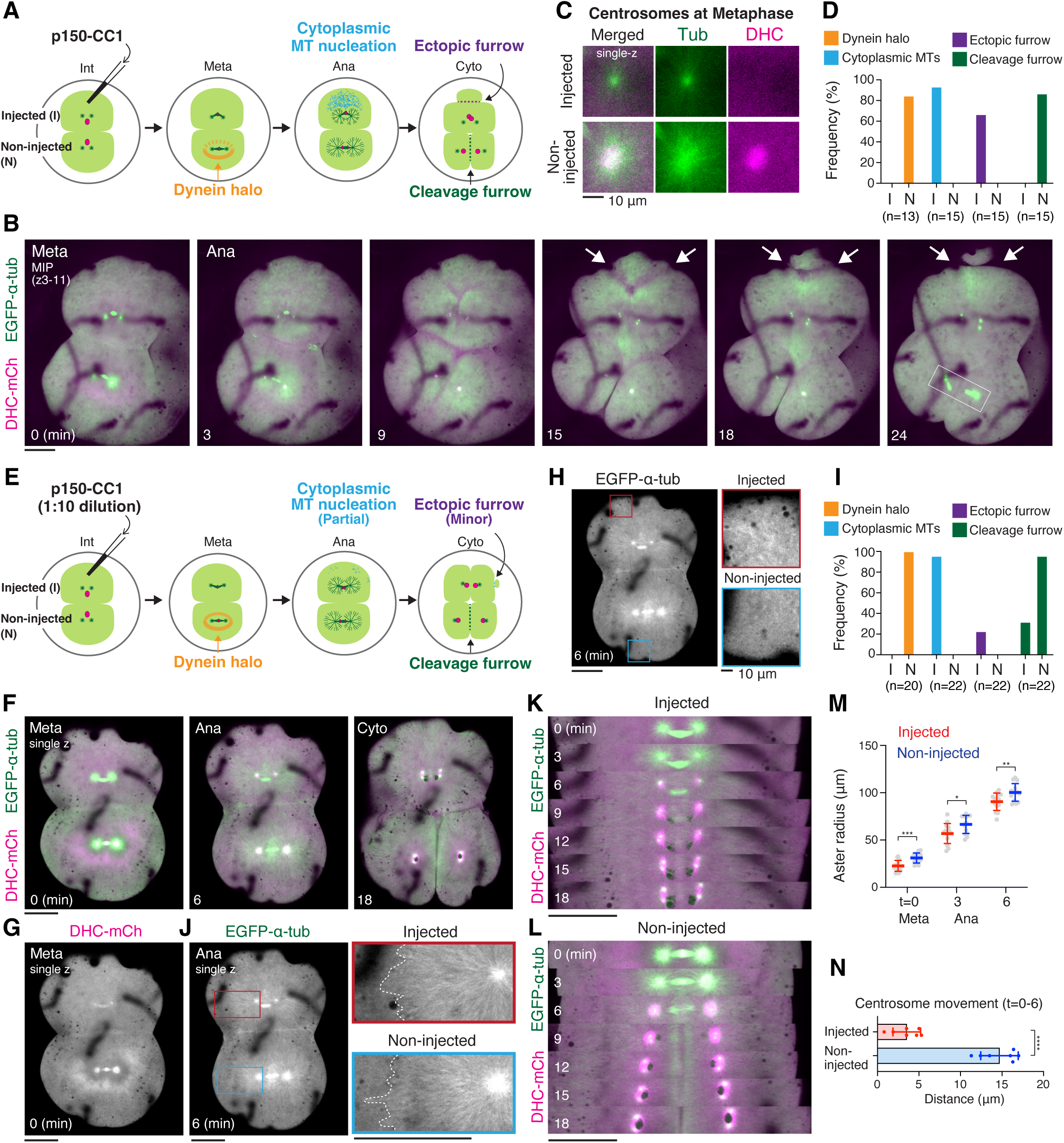
Dynein halo disruption causes cytoplasmic MT nucleation and centrosome positioning defects. (A) Schematics showing phenotypes after p150-CC1 injection in one blastomere of a 2-cell embryo. (B) Live images of MTs (green) and dynein (magenta) in 2-cell blastomeres after p150-CC1 injection. Arrows indicate ectopic furrows. (C) Live fluorescence images of MTs and dynein at metaphase centrosomes in p150-CC1 injected (top) and non-injected (bottom) blastomeres. (D, I) Graphs showing frequencies of indicated phenotypes caused by injection of 20 mg/ml (D) or 2 mg/ml (I) of p150-CC1. Injected (I) and non-injected (N) 2-cell blastomeres were analyzed. (E) Schematics showing phenotypes after injection of 10-fold diluted p150-CC1. (F) Live images of MTs and dynein in 2-cell blastomeres after diluted p150-CC1 injection. (G) A live image showing dynein halo disruption in p150-CC1 injected blastomere (top). (H) Live images of MTs showing cytoplasmic MT nucleation in p150-CC1 injected blastomere (top). (J) Live images of MTs showing anaphase aster growth. (K-L) Kymographs showing MTs (green) and dynein (magenta) in the diluted p150-CC1 injected (top) and non-injected (bottom) blastomeres. (M) Graphs showing aster radii in diluted p150-CC1 injected and non-injected 2-cell blastomeres (n=14). (N) Graphs showing distances of centrosome movement in diluted p150-CC1 injected and non-injected 2-cell blastomeres (n=7). ****p<0.0001. Scale bars = 100 μm except for (C) and (H, right, enlarged) in which scale bars = 10 μm.

To disrupt the dynein halo more specifically, we next injected 10-fold diluted p150-CC1 (Fig. 4E-F). In injected blastomeres, dynein was still detectable at centrosomes and the spindle, but was homogenously distributed in cytoplasm without halo formation (Fig. 4F-G, Movie S8). In anaphase, cytoplasmic MTs nucleated at small regions near the cell periphery in 95.5% (n=21/22) of injected blastomeres (Fig. 4H-I), which caused small bud-like ectopic furrows in 22.7% (n=5/22) of blastomeres (Fig. 4I, Fig. S8G). Intriguingly, aster growth was not obviously attenuated by diluted p150-CC1 injection (Fig. 4J-M). However, outward centrosome movement was significantly impaired (Fig. 4K-L, 4N). These results indicate that the halo-like dynein enrichment is required to generate outward aster/centrosome-moving force in anaphase. In addition, dynein in bulk cytoplasm inhibits ectopic MT nucleation that antagonizes aster growth depending on the amount of abnormal MTs.

### Protein knockdown of dynein or dynactin causes ectopic furrows and centrosome positioning defects

In the above experiments, we used p150-CC1 to inhibit dynein (Fig. 3 and 4). To validate dynein inhibition phenotypes, we next degraded mAID-tagged endogenous dynein (DHC-mACF) or dynactin subunit p50 (p50-mACF) using an auxin-inducible degron 2 (AID2) system (Fig. 5A, S9A)^3,46^. Intensities of endogenous DHC-mACF fluorescence gradually diminished during the 2-cell stage (Fig. 5B, Fig. S9B) and reached ∼10% in 4-cell embryos (Fig. 5B-C). These 4-cell embryos showed monopolar or bent bipolar spindles (Fig. S9C, Movie S9), like p150-CC1-injected embryos (Fig. S8F,Movie S8), leading to embryonic lethality (Fig. 5D). We analyzed mitotic phenotypes in 2-cell embryos, in which bipolar spindles still formed, but the dynein halo was undetectable (Fig. 5E). Like diluted p150-CC1-injected embryos (Fig. 4M-N), asters were still able to grow (Fig. 5F), but centrosome movement was significantly reduced (Fig. 5G). In addition, DHC knockdown caused bud-like ectopic furrows (Fig. 5H-I) coupled with cleavage furrow failure (Fig. 5E), as observed in p150-CC1-injected embryos (Fig. 5B). Similar phenotypes were observed in p50-knockdown embryos (Fig. 5J, Fig. S9D-K). Importantly, p50 knockdown disrupted the halo-like cytoplasmic enrichment of dynein (DHC-mCh) (Fig. 5K), suggesting that not only MTs (Fig. 2C), but also dynactin is required for dynein halo formation. Cytoplasmic MT nucleation was hardly detectable in 2-cell stage embryos, likely due to partial knockdown of AID-fusion proteins (Fig. S9B). In contrast, ectopic MT nucleation was detectable in subsequent 4-cell stage mitosis (Fig. 5L, S9L), in which p50 was more depleted (Fig. S9D). These results confirmed that dynein-dynactin complexes suppress ectopic cytoplasmic MT nucleation and generate aster-pulling forces to ensure proper cytokinesis and centrosome positioning in large embryos.

**Figure 5.**
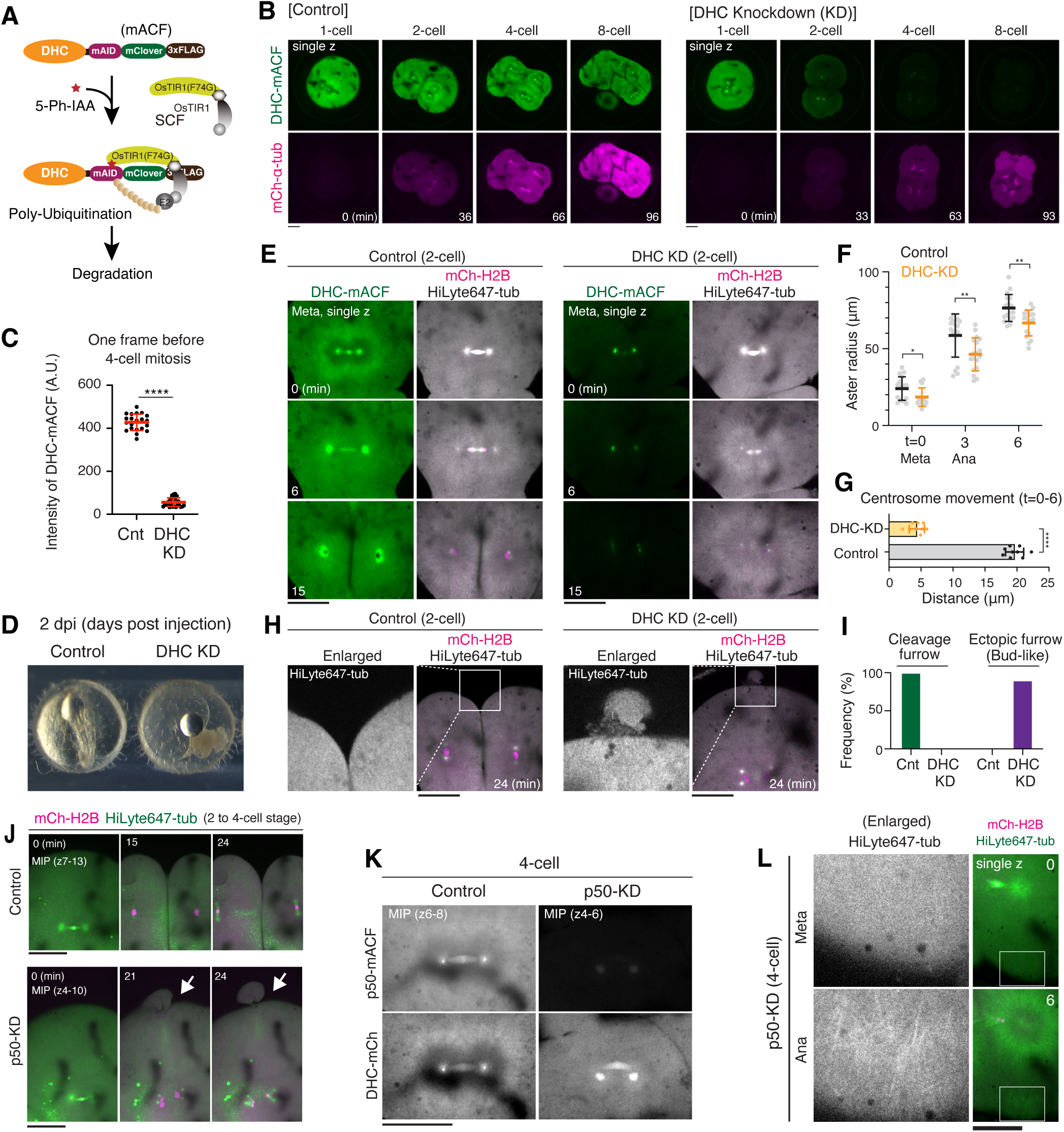
AID-mediated protein knockdown of dynein or dynactin causes centrosome positioning defects and ectopic furrow formation. (A) Schematic representation of auxin-inducible degron 2 (AID2)-mediated DHC degradation. (B) Representative live-cell images showing the fluorescence of DHC-mAID-mClover-3xFLAG (mACF) and mCh-α-tubulin in control (left) and OsTIR1(F74G)-expressing (right) embryos. (C) Quantification of fluorescence intensity of DHC-mACF one frame before 4-cell mitosis in control (n=21) and DHC-knockdown (KD) (n=29) blastomeres. (D) A phase contrast image showing lethality in a DHC-knockdown embryo (right). (E) Live-cell images of control (left) and DHC KD 2-cell blastomeres showing chromosomes (mCh-H2B, magenta) and MTs (HiLyte647-tubulin, white). Fluorescence intensities of endogenous DHC-mACF were adjusted with the same scale between control and DHC-KD blastomeres, but those of MTs and chromosomes were differently adjusted since their fluorescence intensities were variable due to injection amount. (F) Graphs showing aster radii in control (n=17) and DHC-knockdown (n=18) blastomeres. *p<0.1, **p<0.01. (G) Graphs showing distances of centrosome movement in control (n=9) and DHC-knockdown (n=9) blastomeres. ****p<0.0001. (H) Live images showing a normal furrow and a bud-like ectopic furrow in control and DHC-knockdown blastomeres, respectively. (I) Frequencies of a normal cleavage furrow and bud-like ectopic furrows in control (n=10) and DHC-KD (n=10) 2-cell stage embryos. (J) Live images showing a bud-like ectopic furrow in a p50-knockdown blastomere. (K) Live images of p50-mACF and DHC-mCh showing no dynein halo in a p50-knockdown blastomere. mRNAs encoding either miRFP670Nano3-H2B or OsTIR1(F74G)-P2A-miRFP670Nano3-H2B were injected in control or p50-KD embryos, respectively. (L) Live images of MTs (HiLyte647-tubulin) showing anaphase cytoplasmic MT nucleation caused by p50-knockdown in 4-cell blastomeres. Scale bars = 100 μm.

### Localized dynein halo disruption causes asymmetric cytoplasmic MT nucleation and aster positioning

The dynein halo forms symmetrically around the metaphase spindle (Fig. 1B-C). To clarify its function, particularly in generating forces that position centrosomes, we next asymmetrically disrupted the dynein halo using laser ablation. We employed multiple-laser pulses at small region of dynein halo next to one of the two asters (Fig. 6A t= –3), which created asymmetric halo disruption at metaphase (Fig. 6A t=0). As expected, cytoplasmic MTs were nucleated specifically near the ablated region after anaphase onset (Fig. 6B, Fig. S10A-B, Movie S10), occasionally resulting in a pseudo furrow (Fig. S10C). Intriguingly, dynein was detected as weak punctate signals that co-localized with nucleated cytoplasmic MTs (Fig. 6B). Although aster radii were difficult to compare due to potential photo-bleaching of EGFP-α-tubulin by laser ablation, aster movement was significantly impaired by halo disruption (Fig. 6C-D, Fig. S10D).

**Figure 6.**
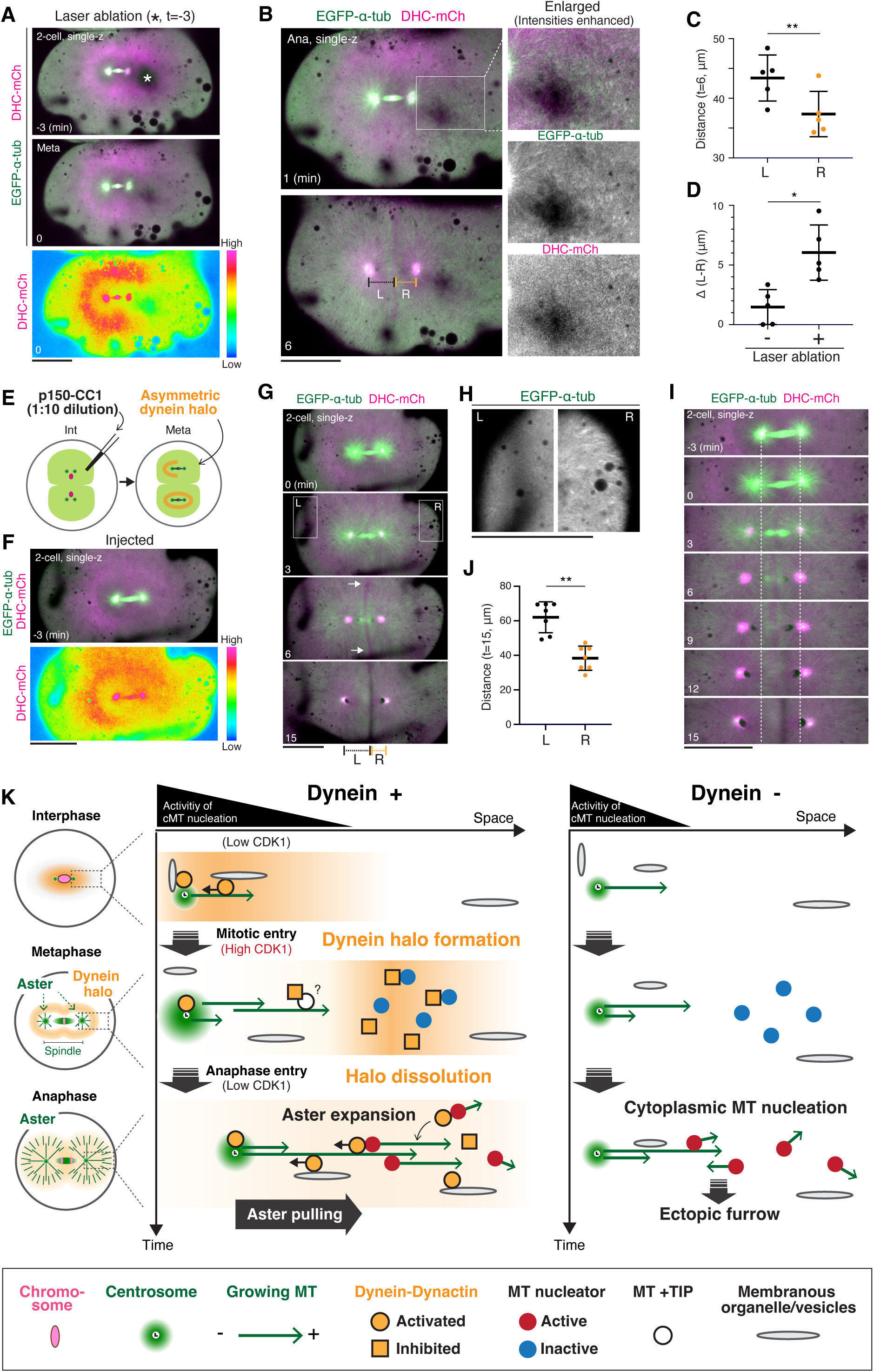
Local disruption of the dynein halo causes asymmetric cytoplasmic MT nucleation and aster positioning defects. (A) Representative images of MTs and dynein with a heatmap image of dynein (bottom) showing local dynein halo disruption by laser ablation. Laser ablation was performed at the region indicated by an asterisk. (B) Live images of MTs and dynein in anaphase (top) and cytokinesis (bottom) showing cytoplasmic MT nucleation and short centrosome movement near the ablation region. (C) Quantification of centrosome movement distance at the left (L) and right (R) side (n=5), **p<0.01 (D) Quantification of the difference (L-R) with or without laser ablation (n=5), *p<0.1 (E) Schematics showing diluted p150-CC1 injection and asymmetric dynein halo formation in the injected 2-cell blastomere. (F) A representative image of MTs and dynein with a heatmap image of dynein (bottom) showing an asymmetric dynein halo created by diluted p150-CC1 injection. (G) Live images of MTs and dynein showing cytoplasmic MT nucleation near the halo-disrupted region (R, t=3), a bent cleavage furrow (arrows, t=6), and asymmetrical centrosome movement (t=15). (H) Enlarged images of MTs in (G). (I) A kymograph showing asymmetrical movement of centrosomes and nuclei (indicated by dark oval regions, t=9-15) in response to asymmetric dynein inhibition. (J) Quantification of distances, L and R, shown in G (n=7), **p<0.01 (K) A model showing cell-cycle-dependent formation and dissolution of the dynein halo, and how the dynein halo facilitates anaphase aster expansion and centrosome positioning in large medaka embryos. Scale bars = 100 μm.

To confirm the above results, we next locally injected diluted p150-CC1 prior to 2-cell mitosis (Fig. 6E). This also created an asymmetric dynein halo during 2-cell metaphase (Fig. 6F), which subsequently caused asymmetric cytoplasmic MT nucleation in anaphase (Fig. 6G-H, Fig. S11A-E). Both asters appeared to grow similarly, but a midzone indicated by the central MT-less region between asters was slightly bent (Fig. 6G t=6) in 77.8% (n=7/9) of blastomeres with asymmetric halo. The bent midzone is likely attributable to impaired aster growth near cytoplasmic MTs or spindle displacement toward the halo-rich side (Fig. 6I t=0-3). The midzone eventually became straight (Fig. 6G, I t=15), but asymmetrically positioned between separating asters (Fig. 6J, Movie S11). In summary, these results confirm that the dynein halo suppresses ectopic MT nucleation and ensures aster growth and pulling for successful furrow positioning.

## Discussion

In this study, we investigated localization and functions of endogenous dynein and dynactin in live medaka early embryos. Unexpectedly, we found that dynein does not distribute uniformly in cytoplasm, but that it concentrates in a halo-like zone at the periphery of short microtubule asters during metaphase (Fig. 1). This localization depends on MTs and dynactin (Fig. 2C, 5K). We propose that upon mitotic entry, most active dynein-dynactin complexes around interphase centrosomes are inactivated downstream of high CDK1 activity, dissociating from their activating adaptors and associated cargos, including ER^17^ (Fig. 6K). These inactive dynein-dynactin complexes are subsequently transported toward the aster periphery by a MT plus-end-binding protein (+TIP) network containing CLIP-170 (Fig. 6K), which interacts with dynactin^47,48^. Transported inactive dynein-dynactin complexes fall off astral MT tips and accumulate as a halo due to the gel-like property of embryonic cytoplasm^49^, and/or confinement by ER or other cytoplasmic compartments (Fig. 2H). After anaphase onset, reduced CDK1 activities likely re-activate dynein at the halo by promoting dynein’s association with its activating adaptors, leading to minus-end-directed transport of cargos on growing asters (Fig. 6K).

Inhibition of dynein using p150-CC1 or genetic protein knockdown had two striking effects, inhibition of aster growth after anaphase onset and promotion of ectopic microtubule nucleation (Fig. 3E, 4B, 5L, S6C). Considering that localized halo disruption by laser ablation also caused ectopic MT nucleation (Fig. 6B), cytoplasmic MT generation is not an artifact, but a phenotype caused by dynein halo inhibition. We propose that the dynein halo normally incorporates these cytoplasmic MTs into anaphase asters with their minus-ends oriented toward centrosomes (Fig. 6K) to facilitate anaphase aster expansion as a “booster” away from centrosomes (Fig. 6K). Similar models of nucleated MT incorporation/transport were proposed for the role of dynein in meiotic spindle assembly^50^. Intriguingly, nucleated cytoplasmic MTs also showed ring-like or semi-ring-like shapes (Fig.3 F-J). It is tempting to speculate that these MT nucleators may also be enriched in the dynein halo in an inactive form at metaphase, and that they are then activated by anaphase initiation (Fig. 6K).

Previous research demonstrated that aster growth is initiated by centrosomes and promoted by MT-dependent autocatalytic MT nucleation^30,31^. The dynein halo would assist this autocatalytic growth mechanism by incorporating MT nucleators or cytoplasmic MTs into asters and/or by suppressing ectopic MT nucleation in anaphase cytoplasm to supply free tubulin to asters (Fig. 6K). Interestingly, partial dynein inhibition or knockdown is insufficient to inhibit aster growth (Fig.4M, 5F), but sufficient to reduce outward centrosome movement (Fig.4N, 5G), indicating that aster growth can be driven by both dynein-dependent and –independent mechanisms^30^, but aster-moving forces are largely generated by dynein. Given that dynein seems to interact with membranous organelles after anaphase onset (Fig. 2C, 2H-I), these findings suggest that dynein localized to bulk cytoplasm, perhaps on organelles, generates outward pulling forces on aster MTs after anaphase (Fig. 6K), as proposed previously^1,22^, especially when cargos are physically connected to or generate friction force by gel-like cytoplasm^15,23,49^.

The dynein halo is a unique cell cycle– and MT-regulated cytoplasmic enrichment of dynein-dynactin complexes (Fig. 6K) that differs from template-based reaction-diffusion gradients such as Ran-GTP^51^. The dynein halo contributes to both self-organization of asters^52^ and centrosome centering in daughter cells^1^, which are required for symmetrical patterning in early vertebrate cleavage^3^. Our understanding of early embryonic division will be further improved by studying detailed mechanisms of dynein-halo formation/dissolution, function, and their adaptation during embryogenesis in various vertebrates.

## Acknowledgments

We thank Toane Arata, Sumika Hagihara, Yoko Nakasone and the OIST animal resource section staffs for feeding and maintenance of medaka fish at OIST. We are grateful to NBRP Medaka (https://shigen.nig.ac.jp/medaka/) for providing OK-Cab (Strain ID: MT830). We thank Yohsuke Moriyama, Kojiro Suda, and Keiko Kono for BiP-mCh-KDEL plasmid, and Christine M. Field for discussion and assistance.

## Funding

This work was supported by grants from JSPS KAKENHI (21H02481, 25K02275, and 25H02403 to TK, and 24K09462 to AK), JST FOREST (JPMJFR224O to TK) and the Takeda Foundation (to TK), the Uehara Foundation (to TK), the Naito Foundation (to TK) and the Okinawa Institute of Science and Technology Graduate University (to TK).

## Author contributions

Conceptualization, TK, LB, and TJM; Investigation, AK; Formal analysis, AK and TK; Methodology, AK and TK; Resources, AK, LB, TJM, and TK; Writing-original draft, TK; Writing-review & editing, TJM, LB, AK, and TK; Supervision, TK; Funding Acquisition, TK and AK.

## Competing interests

The authors declare no competing interests.

## Data and materials availability

All data supporting findings of this study are available in the paper and its Supplementary Materials. All data of this study are stored at the corresponding author and available on reasonable request.

## Supplemental information

Materials and Methods

Figs. S1 to S11

Tables S1 to S4

Movies S1 to S11

## Supplemental information

### Materials and Methods

#### Fish maintenance

Fish experiments were conducted in accordance with protocols (ACUP-2023-009, ACUP-2025-038) approved by the Animal Care and Use Committee at Okinawa Institute of Science and Technology Graduate University (OIST). The OK-Cab strain (MT830) of medaka (*Oryzyas latipes*) was obtained from the National Bio-Resource Project Medaka (NBRP Medaka) and used as the parental strain. Medaka were raised and maintained as described previously^1^.

Naturally fertilized eggs were collected from 2-9 month old pairs. Healthy fertilized eggs were used for imaging. Fluorescence of DHC-mCh, DHC-CF, DHC-mACF, p50-mACF, and RCC1-mCh^1^ were constantly observed in all knock-in embryos. However, fluorescence of EGFP-α-tubulin was variable, as described previously^1^. We selectively used embryos having similar fluorescence intensity of EGFP-α-tubulin in both control and inhibition/depletion experiments. To investigate cytoplasmic MT nucleation, we selected embryos having relatively strong expression of EGFP-α-tubulin.

#### Plasmid and dsDNA construction

Donor plasmids and dsDNAs for CRISPR/Cas9-mediated genome editing (Fig. S1) were designed and constructed as described previously^1^. The designed DNA containing, stop-codon mutation, a BamHI site, and homology arms for Ol-DHC1 or Ol-p50/DCTN2 (450-500 bp homology arms) were synthesized by a gene synthesis service (Genewiz, South Plainsfield, NJ). mCherry2 (mCh), mClover-3xFLAG (CF), or mAID-mClover-3xFLAG(mACF) cassettes were inserted into the BamHI site. dsDNAs were amplified by PCR using modified primers with 5’Bioton – 5 x phosphorothioate bonds^2^ (synthesized by eurofins) and PrimeSTAR Max (Takara). To exogenously express an ER marker, BiP-mCherry-KDEL, cDNA coding BiP-mCherry-KDEL (a kind gift from Yohsuke Moriyama and Keiko Kono, OIST) was cloned into a pCS2+ vector for mRNA synthesis.

Established plasmids, medaka strains, and sequence information about guide RNA (gRNA) and PCR primers used in this study are described in Table S1, S2, S3 and S4, respectively.

#### gRNA synthesis and invitro transcription

For synthesis of gRNA, T7 tagged DNA templates containing the gRNA sequence were amplified by PCR, as described previously^3^. PCR fragments were used as a template for *in vitro* transcription with a MEGAscript T7 kit (Thermo Fisher Scientific, AM1333), according to the manufacturer’s instructions. For synthesis of Cas9 mRNA, the pCS2+hSpCas9 vector (Addgene #51815) was linearized by NotI digestion, followed by *in vitro* transcription using an mMessage mMachine SP6 kit (Thermo Fisher Scientific, AM1340) according to the manufacturer’s instructions. Synthesized RNA was purified with an RNeasy Mini kit (Qiagen). To synthesize other mRNAs, template plasmids were linearized with NotI or BssHII.

#### Microinjection

Glass needles were made from borosilicate glass capillaries (Model No: G100F-4, Warner Instruments) using a needle puller (PC-100, Narishige), and attached to a capillary holder connected to a microinjector Femto Jet 4i (eppendorf). For protein injection, glass needles were made from glass capillaries (Narishige, GD-1) coated with Sigmacote (Sigma, SL2-25ML). Needles were manually controlled using a micro-manipulator (MN-153, Narishige) on the stage of a stereomicroscope (Leica M80).

To exogenously express EGFP– or mCherry-fusion proteins, 150 ng/µL mRNAs were injected into one-cell embryos. To visualize MTs simultaneously with green and red fluorescent proteins in early embryos, HiLyte 647-tubulin protein (Cytoskeleton, Inc. TL670M) was injected. 20 μg HiLyte 647-tubulin were resuspended in 5 μL of RNase-free water. After centrifugation (14,000 x g, 4°C, 10 min), supernatant was collected, and aliquots were frozen in liquid nitrogen and stored at –80°C. To inject HiLyte 647-tubulin and mRNAs, the mixture with final concentrations of 800 ng/μL tubulin and 150 ng/µL mRNA was injected using siliconized glass-needles.

Chicken p150-CC1 fragment (pVEX-CC1)^4^ was purified as described previously^5^. 20 mg/mL p150-CC1 was injected in Fig. 3 and Figs. 4A-D, whereas 10-fold diluted p150-CC1 (2mg/mL) was used in Figs. 4E-N and Figs. 6E-I. Since injected embryos must be set up quickly for microscope observation, non-injected embryos served as controls. Injection of p150-CC1 dialysis buffer (XB buffer: 100 mM KCl, 10 mM HEPES pH7.7, 1 mM MgCl_2_, 1 mM EGTA) in one-cell zygotes (n=12) or 2-cell stage embryos (n=21) did not cause cytoplasmic MT nucleation and ectopic furrows.

#### Establishment of knock-in strains

To generate DHC-mCh, DHC-CF, DHC-mACF or p50-mACF knock-in strains (Fig. S1), a mixture of 50 ng/µL gRNAs, 150 ng/µL Cas9 mRNA and 10 ng/µL dsDNAs was injected into one-cell stage medaka embryos. After rearing injected embryos to adult fish, small pieces of tail fin were used for genomic PCR with KOD Fx Neo (TOYOBO) as described previously^1^ using primers listed in Table S4. Candidate G0 fish were crossed with wild-type counterparts, and fertilized eggs were analyzed for fluorescence using a fluorescence stereo microscope (Leica M205 FCA) 1 or 2 days after fertilization. Genomic DNA was extracted from some fluorescence-positive F1 embryos, and genomic PCR was performed. Proper knock-in of the construct was confirmed by direct sequencing of PCR products. Other positive embryos from the same G0 fish were reared, and F1 male and female fish were mated to obtain a homozygous F2 generation. PCR products were analyzed by normal gel electrophoresis (Fig. S1C) or using an automatic microchip electrophoresis system (MCE-202 MultiNA: Shimazu, Kyoto, Japan) (Figs. S1A, B, D).

#### Microscope system and live imaging

Imaging was performed with spinning-disc confocal microscopy with a 20× 0.95 numerical aperture objective lens (APO LWD 20X WI λS, Nikon, Tokyo, Japan). A CSU-W1 confocal unit (Yokogawa Electric Corporation, Tokyo, Japan) with three lasers (488, 561, 640 nm, Coherent, Santa Clara, CA) and an ORCA-Fusion digital CMOS camera (Hamamatsu Photonics, Hamamatsu City, Japan) were attached to an ECLIPSE Ti2-E inverted microscope (Nikon) with a perfect focus and a water-supply system. Laser power of 488, 561, and 640 nm lasers was measured using a NovaII power meter (Ophir) and a PD300-MS microscope slide power sensor (Ophir), and adjusted to 2.90, 3.50, and 2.38 mW, respectively, at the objective lens. Images were captured using NIS-Elements software (version 5.21.00, Nikon). Samples were imaged at room temperature (24-26°C).

To hold living embryos for time-lapse imaging, a handmade device was created: 1.5 % agarose solution was added to glass-bottomed dishes (CELLview™, #627860, Greiner Bio-One, Kremsmünster, Austria), and a 7-unified cover-glass (18 mm x 18 mm with 0.12-0.17 mm in thickness, Matsunami) or custom-made mold was set in the center of the glass-bottomed dish. When the agarose became solid, the unified glass or mold was removed to make a concave pocket in the agarose gel, 0.8-1.2 mm in width and 3-5 mm in height on the bottom glass (Fig. S9A). 2-8 embryos were put in the pocket in a row with their blastodiscs facing the glass (Fig. 1A). The dish was filled with ∼2 mL medaka balanced salt solution (BSS, 0.65% NaCl, 0.04% KCl, 0.02% MgSO_4_・7H_2_O, 0.02% CaCl_2_・2H_2_O, sterilized and adjusted to pH 8.3 with 5% NaHCO_3_) without phenol red. 11-13 z-section images with a 5 μm z-step were acquired every 3 min, 1 min, or 30 sec with camera binning 1. Green, red, and far-red fluorescence images were captured with exposure times of 500 msec, 1 sec, and 300 msec, respectively. X-Y-Z positions of 2-7 embryos were memorized and automatically imaged in a time-lapse experiment. For figures, 8-bit, maximally projected z-stack images (MIP) or single z-section images are shown as indicated. MIP images were created using NIS-Elements. Signals were linearly adjusted using NIS-Elements and Photoshop to optimize image clarity. Final images were arranged using Adobe Illustrator. Phase-contrast images of embryos for Fig. 5D and S9F were taken with a Leica M80 stereo microscope and a Leica MC190 HD camera.

For nocodazole treatment (Fig. 2C), embryos were incubated in a 24-well plate containing 170-nM nocodazole (Sigma-Aldrich, M1404) in BSS for 10-30 min at 27.5°C. Then, eggs were moved into a hand-made agarose pocket on the glass-bottom dish containing BSS with nocodazole. In Fig. 2I, injected embryos were treated with 2 μM nocodazole to more efficiently depolymerize MTs during 4-cell mitosis.

#### AID2-mediated protein knockdown

For AID2-mediated protein degradation, mRNAs encoding OsTIR1(F74G)-P2A-mCherry-α-tubulin, OsTIR1(F74G)-P2A-mCherry-H2B, or OsTIR1(F74G)-P2A-miRFP670Nano3-H2B were injected into one-cell embryos (Fig. S9A). Corresponding constructs without OsTIR1(F74G) were injected as controls, which normally generated fluorescent proteins more efficiently.

Injected embryos were cultured in BSS without phenol red for 10-30 min with gentle shaking. BSS was replaced with 2 mL BSS containing 10 mM 5-Ph-IAA or 0.1% DMSO, and then these embryos were transferred with the solution to the handmade glass-bottom dish (Fig. S9A). After adjusting the orientation of eggs in the agarose pocket, eggs were observed using the inverted-microscope.

#### Laser ablation

Laser ablation was performed using the MicroPoint Laser System (Photonic Instruments Co., Ltd) with a cartridge-type laser unit. The wavelength of the Dye Laser was 435 nm, and the pulse width (FWHM, full width a half maximum) was 3.5 ns.

#### Quantification

NIS-Elements software (version 5.41.00) was used for line scans. The line width is indicated in figure legends and the mean intensities are shown in the graph. For quantification of aster radii, single focal z-section images with aster centers were selected, and length from the aster center to the aster tip was manually measured using NIS-Elements software. The clearly measurable aster radii were quantified for 3 arbitrary orientations, roughly two vertical and one horizontal orientation to the spindle axis, and the average length was calculated for each aster. Distance of centrosome movement (D) during the indicated duration (t=A-B) was determined by the following formula: D = [Centrosome-centrosome distance (t=B) – Centrosome-centrosome distance (t=A)] x 1/2. Mean fluorescent intensities of DHC-mACF or p50-mACF in the cytoplasm were measured one frame before NEBD at the 4-cell mitosis using NIS-Elements, and neighboring background fluorescence intensities were subtracted in Fig. 5C and Fig. S9E.

#### Illustration and Supplementary movies

Diagrams were created using Adobe Illustrator 2025 (version 29.8.1, Adobe). Supplementary movies were created using NIS-Elements, Fiji, and Adobe Express. To reduce file size, the resolution of movies S2 and S9 was lowered.

#### Statistics and Reproducibility

GraphPad Prism (version 10.2.3) and Excel were used. Mean, SD, and statistical significance were determined using two-sided Welch’s t-tests (Figs.3K, 3L,4M, 4N, 5C, 5F, 5G, 6C, 6D, 6J and Supplementary Figs. S8D, S8E, S9E, S9H, S9I). P values are shown as *: *: p<0.1, **: p<0.01, ***: p<0.001 and ****: p<0.0001. Healthy fertilized eggs were used randomly. At least 3 independent experiments were performed, and similar results were obtained in repeated experiments. No statistical method was used to predetermine sample size. Sample sizes were chosen based on standards in the field and are sufficient for appropriate statistical tests. Investigators were not blinded for data collection and quantification.

**Table S1.** Plasmids used in this study.

| No | Name | Description | Figures | Source |
| --- | --- | --- | --- | --- |
| 1 | pTK938 | dsDNA template having mCherry2 CDS and OI-DHC/dync1h1 homology arms | Fig. S1A | This study |
| 2 | pTK936 | dsDNA template having mClover-3xFLAG CDS and OI-DHC/dync1h1 homology arms | Fig. S1B | This study |
| 3 | pTK1021 | dsDNA template having mAID-mClover-3xFLAG CDS and OI-DHC/dync1h1 homology arms | Fig. S1C | This study |
| 4 | pTK1092 | dsDNA template having mAID-mClover-3xFLAG CDS and OI-p50/DCTN2 homology arms | Fig. S1D | This study |
|  | Addgene #51815 | pCS2+hSpCas9 | Figs. S1A-D | <sup>6</sup> |
| 5 | pTK1095 | pCS2+EMTB-3xGFP | Figs. 1G, 1H, S3A, S3B | <sup>1</sup> |
| 6 | pTK1113 | pCS2+BiP-mCh-KDEL | Figs. 2H, 2I, S5B | This study |
| 7 | pTK1073 | pCS2+mCh-α-tubulin | Figs. 5B, S9C, S9D | <sup>1</sup> |
| 8 | pTK1076 | pCS2+mCh-H2B | Figs. 5E, 5H, 5J, 5L, S9G, S9K, S9L | <sup>1</sup> |
|  | pTK1112 | pCS2+OsTIR1(F74G)-P2A-mCh-H2B | Figs. 5E, 5H, 5J, 5L, S9G, S9K, S9L | <sup>1</sup> |
| 9 | pTK1026 | pCS2+OsTIR1(F74G)-P2A-mCh-α-tubulin | Figs. 5B, S9C, S9D | <sup>1</sup> |
| 10 | pTK1063 | pCS2+miRFP670Nano3-H2B | Fig. 5K | <sup>1</sup> |
| 11 | pTK1111 | pCS2+OsTIR1(F74G)-P2A- miRFP670Nano3-H2B | Fig. 5K | This study |

**Table S2.**
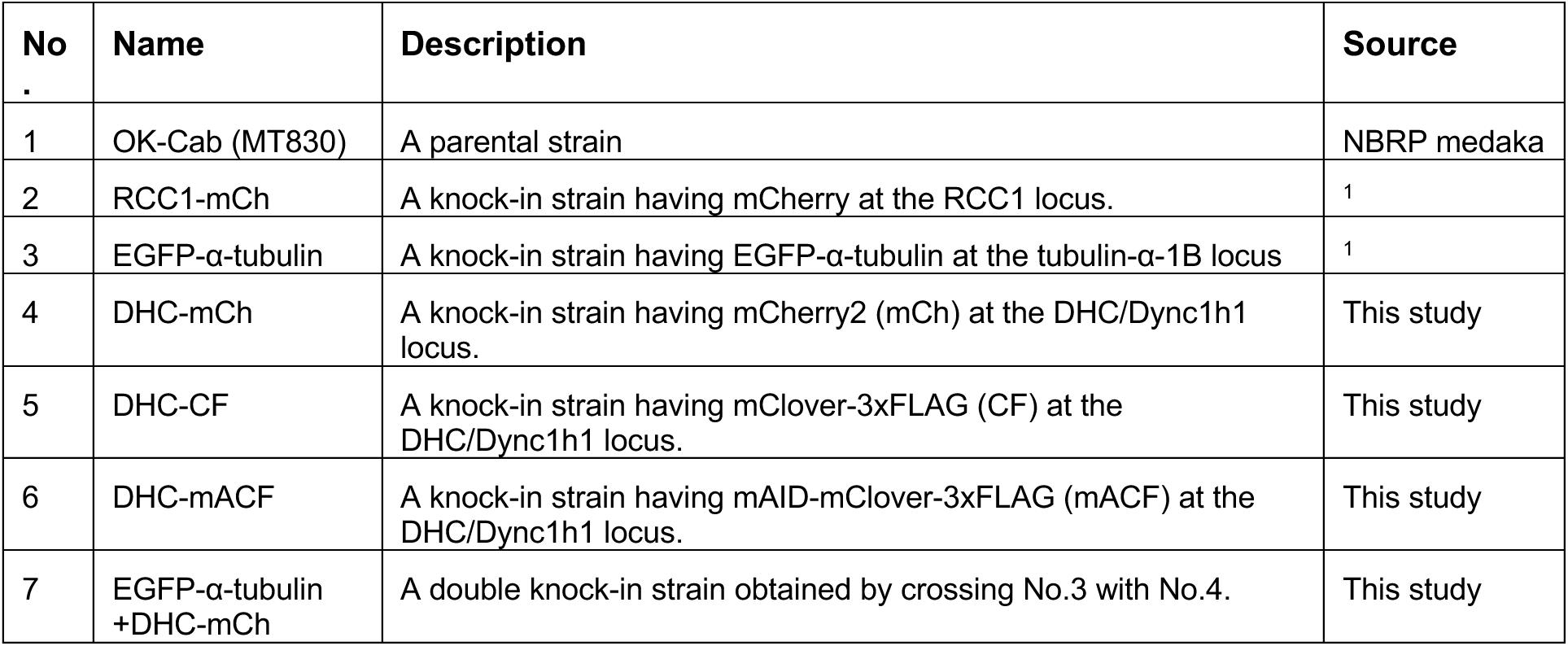
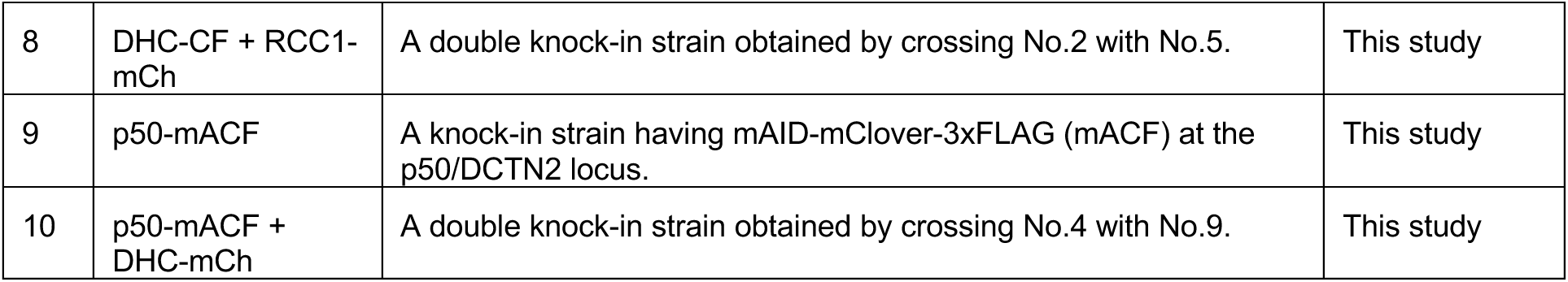
Medaka strains used in this study.

| No | Name | Description | Source |
| --- | --- | --- | --- |
| 1 | OK-Cab (MT830) | A parental strain | NBRP medaka |
| 2 | RCC1-mCh | A knock-in strain having mCherry at the RCC1 locus. | <sup>1</sup> |
| 3 | EGFP-α-tubulin | A knock-in strain having EGFP-α-tubulin at the tubulin-α-1B locus | <sup>1</sup> |
| 4 | DHC-mCh | A knock-in strain having mCherry2 (mCh) at the DHC/Dync1h1 locus. | This study |
| 5 | DHC-CF | A knock-in strain having mClover-3xFLAG (CF) at the DHC/Dync1h1 locus. | This study |
| 6 | DHC-mACF | A knock-in strain having mAID-mClover-3xFLAG (mACF) at the DHC/Dync1h1 locus. | This study |
| 7 | EGFP-α-tubulin +DHC-mCh | A double knock-in strain obtained by crossing No.3 with No.4. | This study |
| 8 | DHC-CF + RCC1-mCh | A double knock-in strain obtained by crossing No.2 with No.5. | This study |
| 9 | p50-mACF | A knock-in strain having mAID-mClover-3xFLAG (mACF) at the p50/DCTN2 locus. | This study |
| 10 | p50-mACF + DHC-mCh | A double knock-in strain obtained by crossing No.4 with No.9. | This study |

**Table S3.**
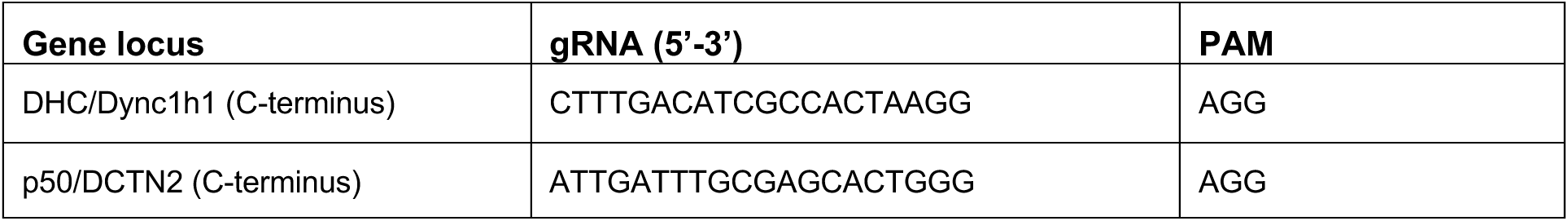
gRNA sequences for CRISPR/Cas9-mediated genome editing.

**Table S4.**
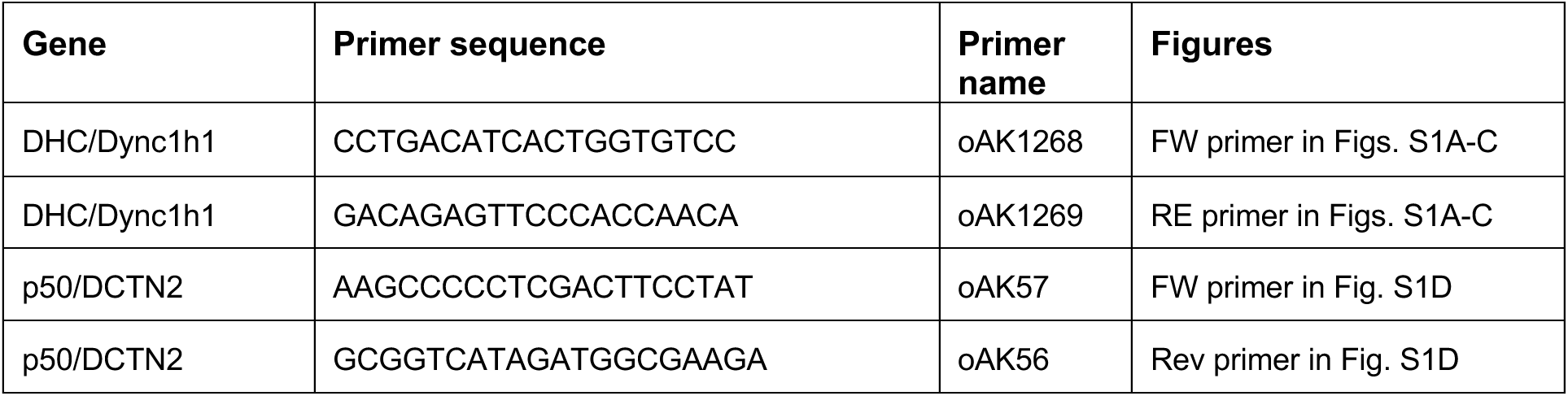
PCR primers used to confirm gene editing.

**Movie S1, related to Fig. 1B**. Time-lapse movies showing MTs (GFP-α-tubulin, green) and endogenous dynein (DHC-mCh, magenta) from 1-cell to 4-cell embryos. MIP images with 6 z-sections are shown.

**Movie S2, related to Fig. 1B**. Grayscale time-lapse movies showing MTs (GFP-α-tubulin, left) and endogenous dynein (DHC-mCh, right) from 1-cell to 4-cell embryos. MIP images with 6 z-sections are shown.

**Movie S3, related to Fig. 1G**. Time-lapse movies showing endogenous dynein (DHC-mCh, magenta) and MTs (EMTB-3xGFP, green) in a 2-cell stage blastomere. Single-z images are shown. Fluorescent intensities of EMTB-3xGFP gradually increase in response to translation of injected mRNAs encoding EMTB-3xGFP.

**Movie S4, related to Fig. 2E**. Time-lapse movies showing endogenous dynein (DHC-mCh, magenta) and MTs (GFP-α-tubulin, green) in a 2-cell stage embryo after centrosome disruption by laser ablation. Single-z images are shown.

**Movie S5, related to Fig. 3E**. Time-lapse movies showing endogenous dynein (DHC-mCh, magenta) and MTs (GFP-α-tubulin, green) in control (left) and p150-CC1 injected (right) zygotes. Single-z images are shown.

**Movie S6, related to Fig. S6C.** Grayscale time-lapse movies showing MTs (GFP-α-tubulin) in control (left) and p150-CC1 injected (right) zygotes. Single-z images are shown.

**Movie S7, related to Fig. 4B**. Time-lapse movies showing endogenous dynein (DHC-mCh, magenta) and MTs (GFP-α-tubulin) in a 2-cell stage embryo. p150-CC1 is injected into one of the blastomeres (top). Single-z images are shown.

**Movie S8, related to Fig. 4F**. Time-lapse movies showing endogenous dynein (DHC-mCh, magenta) and MTs (GFP-α-tubulin) in a 2-cell stage embryo. 10-fold diluted p150-CC1 is injected into one of the blastomeres (top). Single-z images are shown.

**Movie S9, related to Fig. 5B**. Time-lapse movies showing endogenous dynein (DHC-mACF, green) and exogenously expressed mCh-α-tubulin (magenta) in control (–OsTIR1F74G, left) and dynein knockdown (+OsTIR1F74G, right) embryos. The bottom images show only DHC-mACF (green). MIP images with 10 z-sections are shown.

**Movie S10, related to Fig. 6A**. Time-lapse movies showing endogenous dynein (DHC-mCh, magenta) and MTs (GFP-α-tubulin, green) in a 2-cell stage blastomere after local dynein-halo disruption by laser ablation. MIP images with 4 z-sections are shown.

**Movie S11, related to Fig. 6G**. Time-lapse movies showing endogenous dynein (DHC-mCh, magenta) and MTs (GFP-α-tubulin, green) in a 2-cell stage blastomere after asymmetrical dynein-halo disruption by diluted p150-CC1 injection. MIP images with 3 z-sections are shown.

---

**Figure S1.**
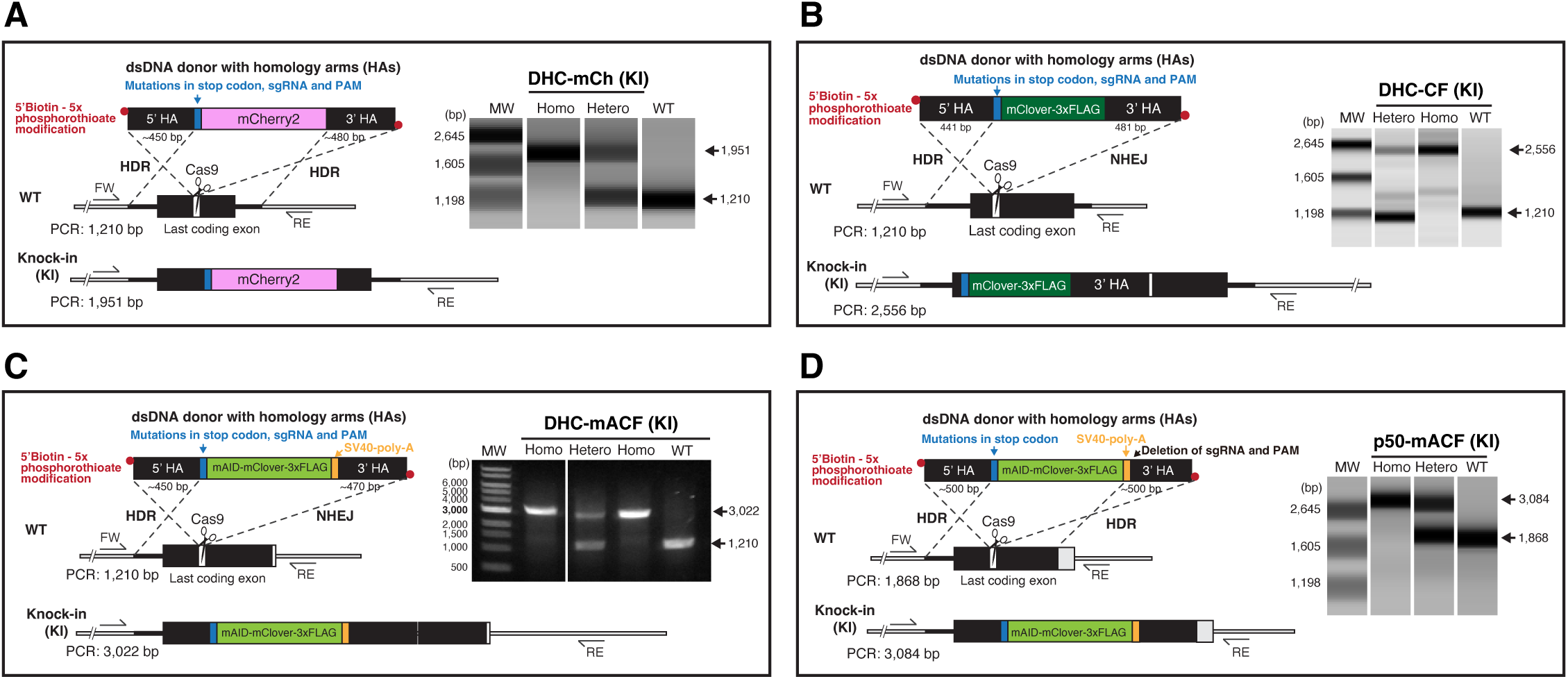
Establishment of knock-in strains for dynein and dynactin. (A-D) Left: schematic representation of generation of the DHC-mCh (A), DHC-CF (B), DHC-mACF (C) or p50-mACF (D) knock-in (KI) strain using a dsDNA donor. Sequence analyses of PCR fragments confirmed that both 5’ and 3’ homology arms (HAs) of the dsDNA donor for DHC-mCh (A) and p50-mACF (D) were integrated into the genome via homology-directed repair (HDR), whereas 3’ HAs of dsDNA for DHC-CF (B) and DHC-mACF (C) were inserted by non-homologous end joining (NHEJ). Right: PCR-based genotyping of the DHC (A-C) or p50 (D) gene in the parental wild-type (WT) and KI strains.

**Figure S2.**
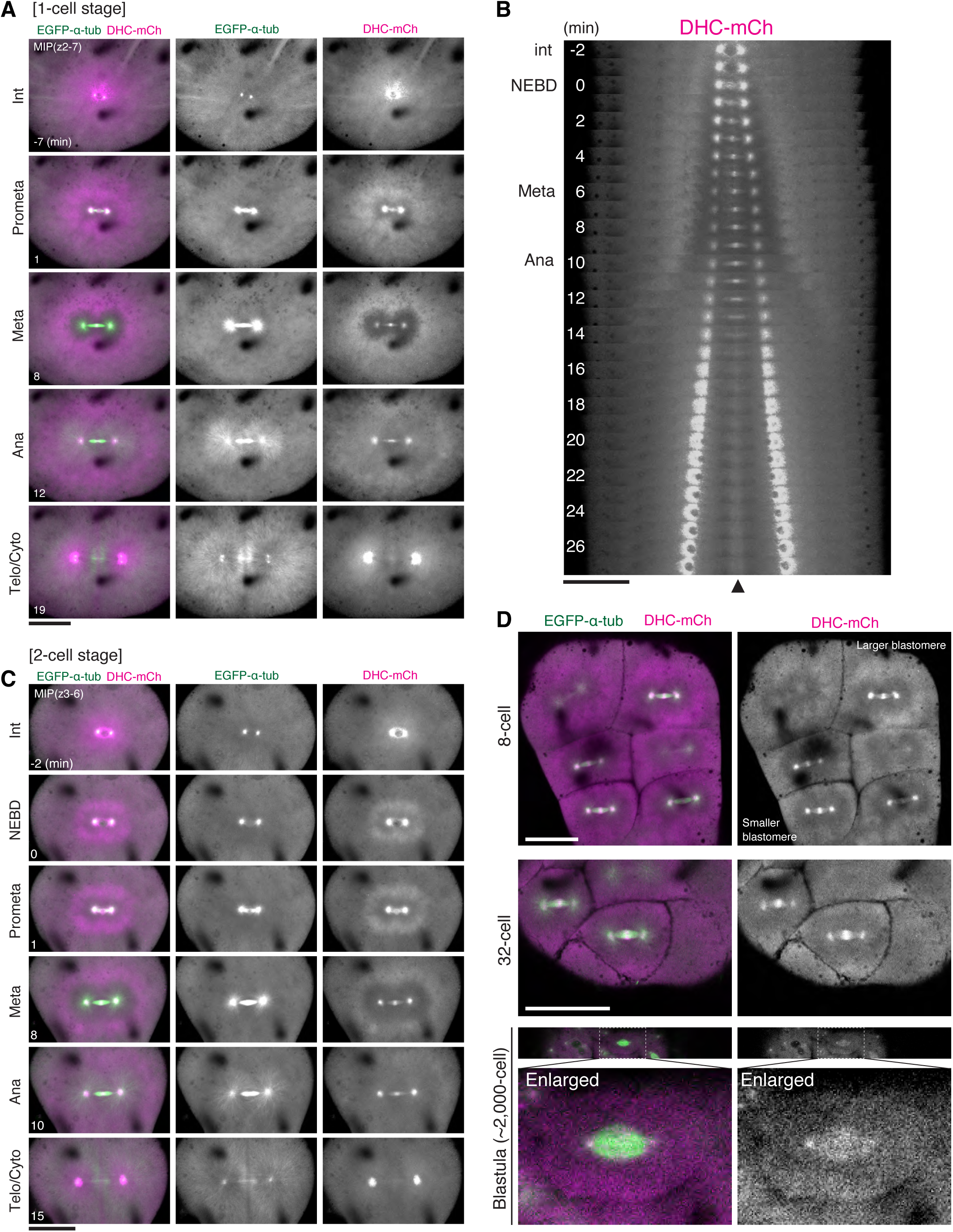
Dynein halo formation at the 1-cell, 2-cell, and later embryonic stages. (A, C) Maximum intensity projection (MIP) images of a live embryo showing localization dynamics of MTs (EGFP-α-tubulin, green) and dynein (DHC-mCh, magenta) during the 1-cell (A) and 2-cell stage (C) mitosis. (B) A kymograph showing dynamic localization change of dynein (DHC-mCh, magenta) during the first mitosis. (D) Live single z-section images of the 8-cell, 32-cell and blastula stage embryos. The dynein halo reached the cell cortex in smaller blastomeres after the 8-cell stage, although dynein appeared to be excluded near the spindle throughout the early embryogenesis. Scale bars = 100 μm.

**Figure S3.**
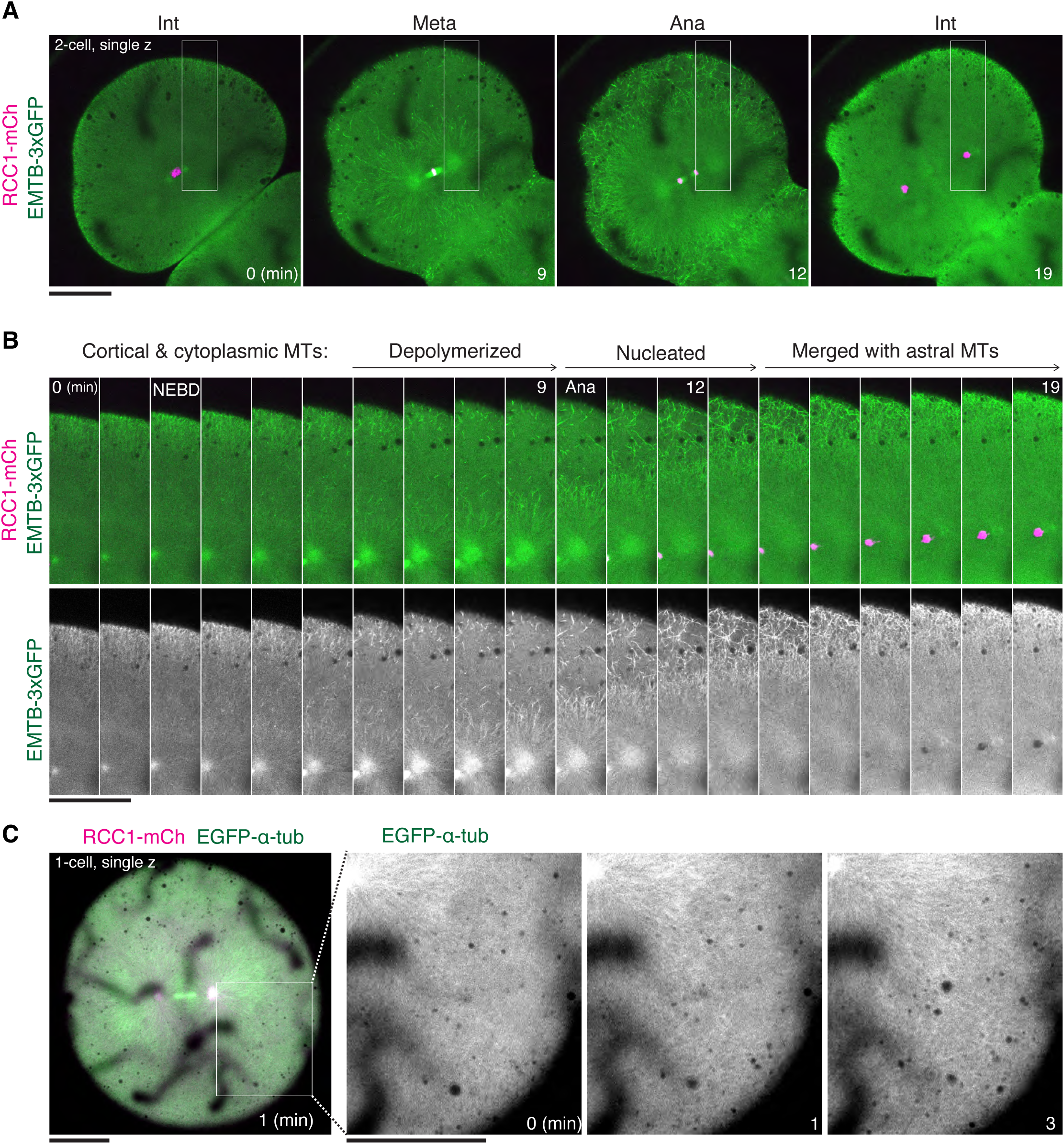
Dynamic behaviors of anaphase asters and cytoplasmic MTs visualized using EMTB-3xGFP or EGFP-α-tubulin in medaka early embryos. (A) Live images showing localization of chromosomes (RCC1-mCh) and asters visualized by EMTB-3xGFP in a 2-cell stage blastomere. Regions indicated by white rectangles were enlarged and used for the kymograph in (B). (B) A kymograph showing the dynamic behaviors of cytoplasmic MTs and asters visualized by EMTB-3xGFP. Interphase cortical and cytoplasmic MTs were depolymerized during mitosis. After anaphase, a subset of MTs appeared to be nucleated in cytoplasm, but they were eventually merged with the growing asters. (C) Left: a single z-section image of a zygote after anaphase. Right: enlarged images showing that cytoplasmic MT nucleation can be visualized by EGFP-α-tubulin, although they are rarely detectable. Scale bars = 100 μm.

**Figure S4.**
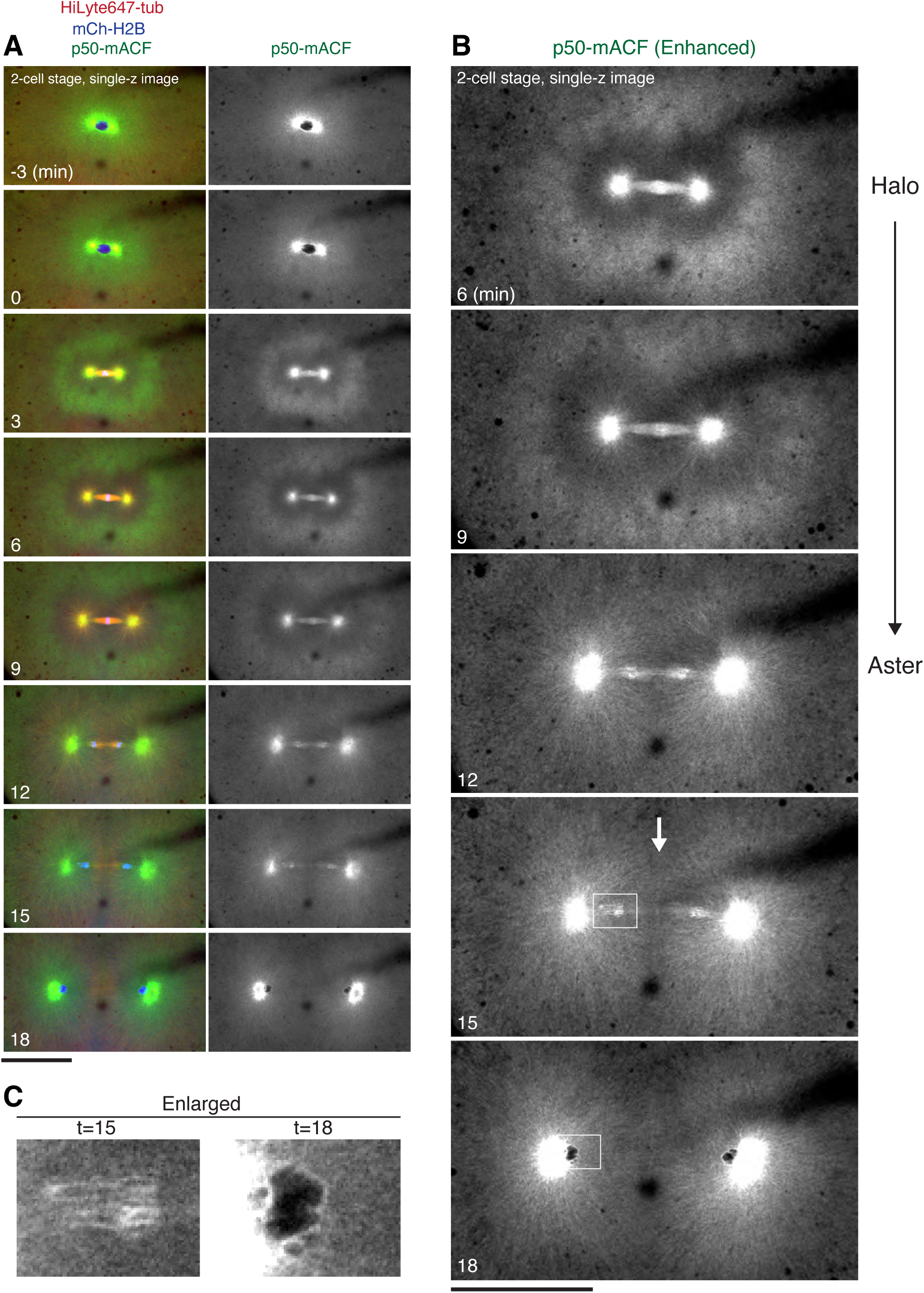
Cell-cycle dependent localization change of endogenous p50 in medaka early embryonic division. (A) Single z-section images of a live 2-cell stage blastomere showing cell-cycle-dependent localization dynamics of dynactin (p50-mACF, green) together with MTs (HiLyte647-tubulin, red) and chromosomes (mCh-H2B, blue) during mitosis. (B) Enlarged, fluorescence-enhanced images of p50-mACF from metaphase to anaphase showing dynamic relocation of p50 from the halo-like cytoplasmic concentration (t=6) to anaphase asters (t=12). p50-mACF was also detectable at the cleavage furrow (arrow, t=15), and at the nuclear envelope of karyomeres (t=18). (C) Enlarged images of p50-mACF around the nuclear envelope of karyomeres. Scale bars = 100 μm.

**Figure S5.**
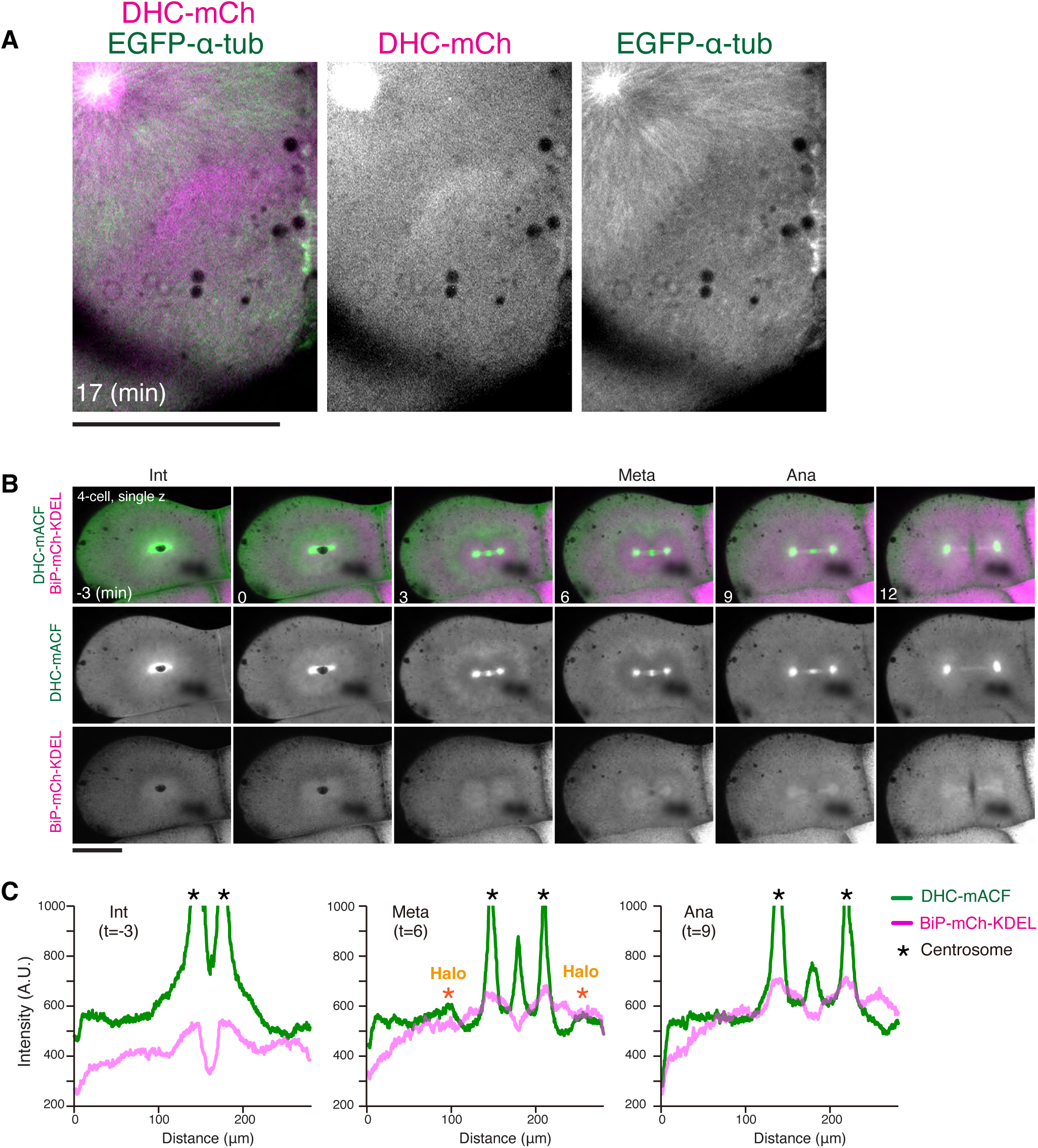
Localization change of dynein and ER during early embryonic division. (A) Enlarged images of Fig. 4E (t=17) showing the remaining dynein halo after anaphase onset. Cytoplasmic MT nucleation was also observed away from the centrosome. (B) Sequential time-lapse images of dynein (DHC-mACF) and an ER marker (BiP-mCh-KDEL) shown in Fig. 2H. (C) Graphs of fluorescence intensities for line scans (line width 32 px) of interphase (t=-3), metaphase (t=6), and anaphase (t=9) blastomeres in (B). Intensities of ER are reduced on the dynein halo at metaphase. Intensities of ER gradually increased due to the translation of injected mRNAs encoding the ER marker. Scale bars = 100 μm.

**Figure S6.**
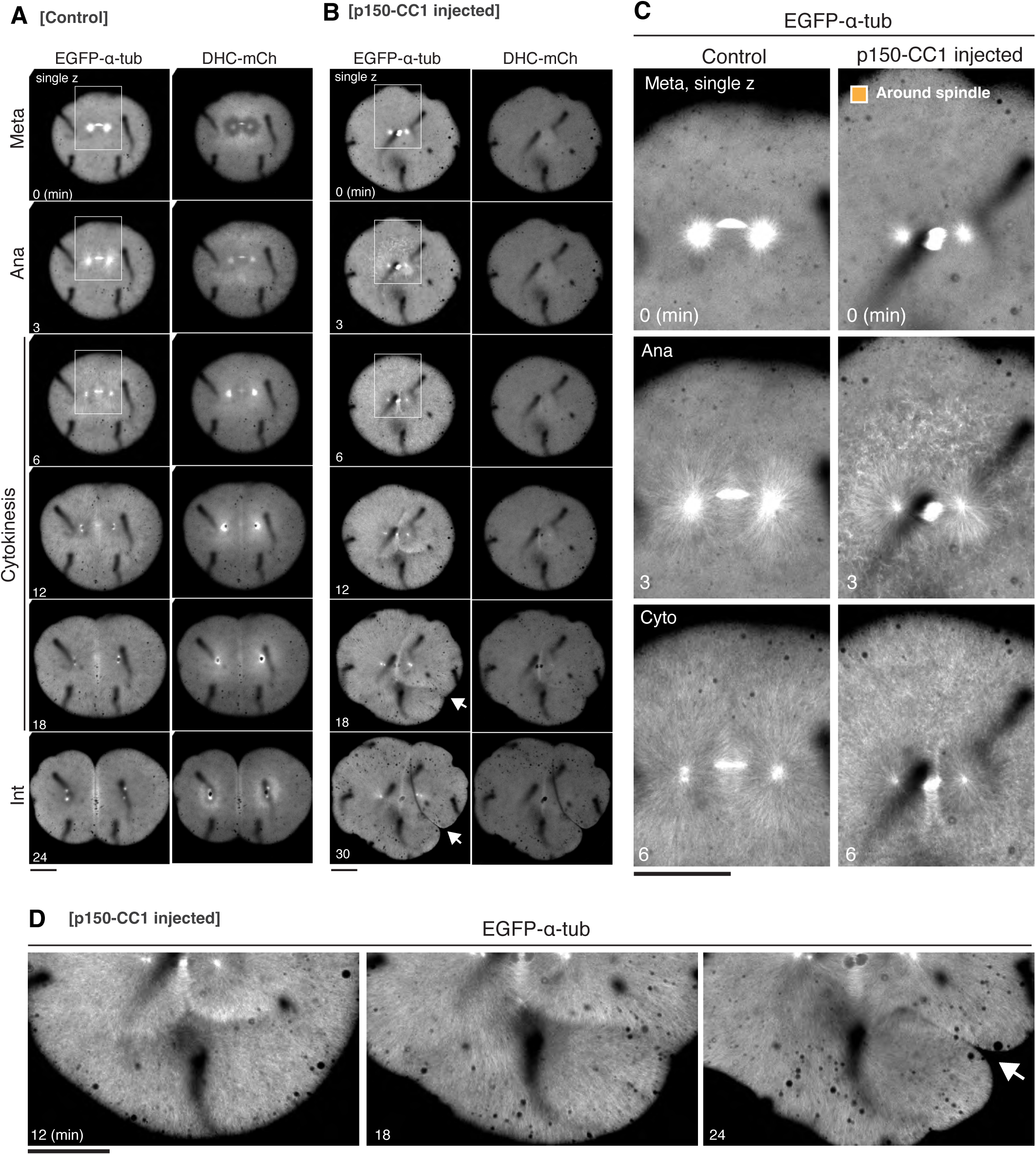
Cytoplasmic MT nucleation in anaphase leads to ectopic furrow formation. (A, B) Another set of control (A) and p150-CC1 injected (B) embryos showing MTs (EGFP-α-tubulin) and endogenous dynein (DHC-mCh). Intensities of MTs and dynein are adjusted equally between control and p150-CC1 injected embryos. Arrows indicate an ectopic furrow. (C) Enlarged images indicated by squares in (A) and (B). p150-CC1 injected embryo showed cytoplasmic MT nucleation around the spindle in anaphase (t=3), which appeared throughout cytoplasm in the next time frame (t=6). (D) Enlarged images of Fig. 3M showing how cytoplasmic MTs merged with asters or created an ectopic furrow (arrow, t=24) at the boundary between cytoplasmic MTs and the aster. Scale bars = 100 μm.

**Figure S7.**
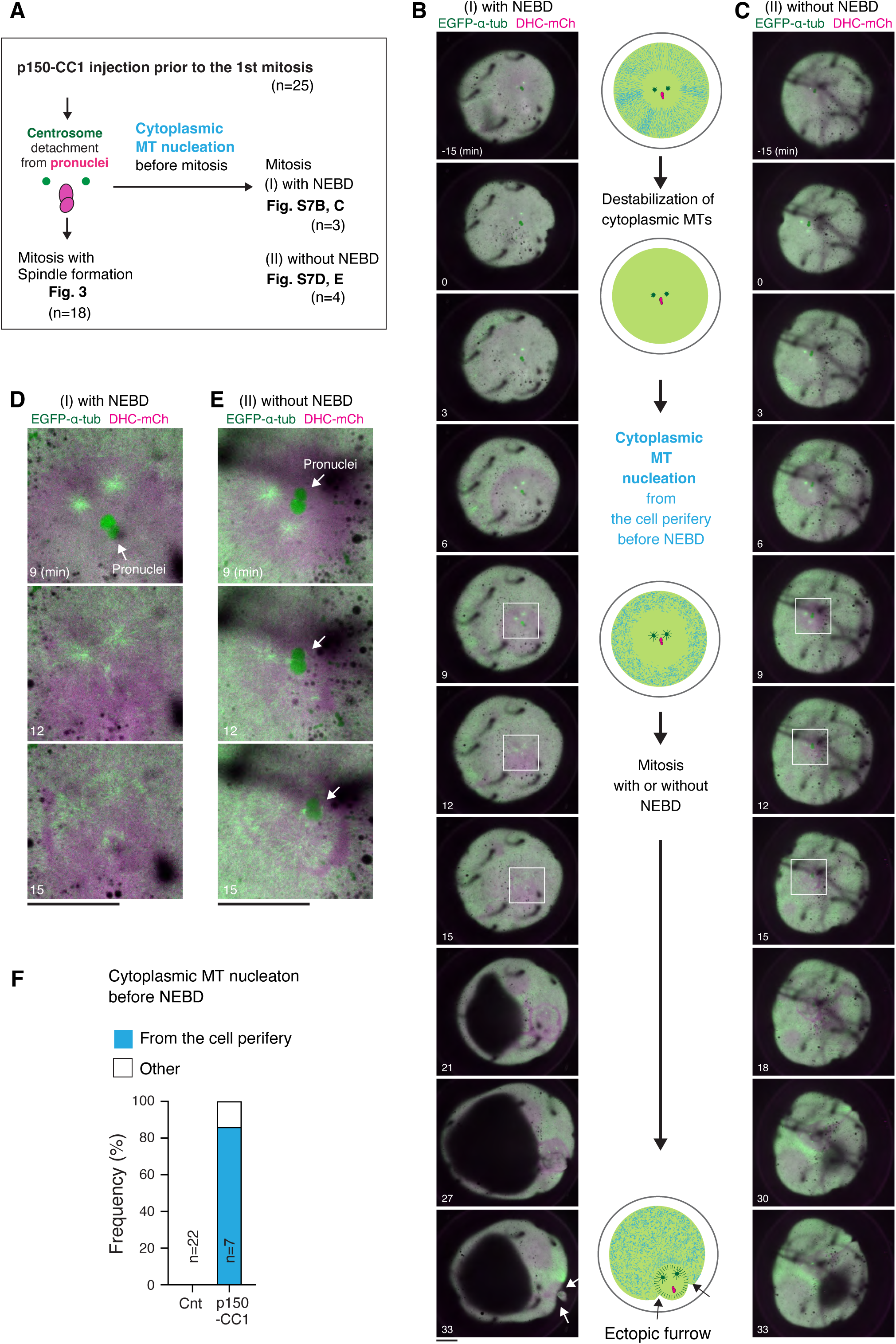
Cytoplasmic MT nucleation from the cell periphery caused by p150-CC1 injection in medaka zygotes. (A) Schematic diagram showing classification of phenotypes caused by p150-CC1 injection prior to the first mitosis. 72% of injected embryos (n=18/25) entered mitosis and formed zygotic spindles, whereas the remaining 18% (n=7/25) showed cytoplasmic MT nucleation before NEBD. (B, C) Single z-section images showing cytoplasmic MT nucleation from the cell periphery to the cell center before the NEBD. The pronuclei characterized by EGFP-α-tubulin enrichment and DHC-mCh exclusion appeared to be broken during mitotic progression (t=12) in 3 out of 7 embryos (B), whereas the pronuclei was not broken in the remaining 4 zygotes (C). (D, E) Enlarged images of rectangular regions in B (D) and C (E) showing mitotic progression with or without the NEBD, respectively. (F) Graphs showing frequencies of cytoplasmic MT nucleation before NEBD in control (n=22) and p150-CC1 injected zygotes (n=7). In most p150-CC1 injected embryos (n=6/7), cytoplasmic MTs nucleated around the cell periphery, and then spread toward the cell center. Scale bars = 100 μm.

**Figure S8.**
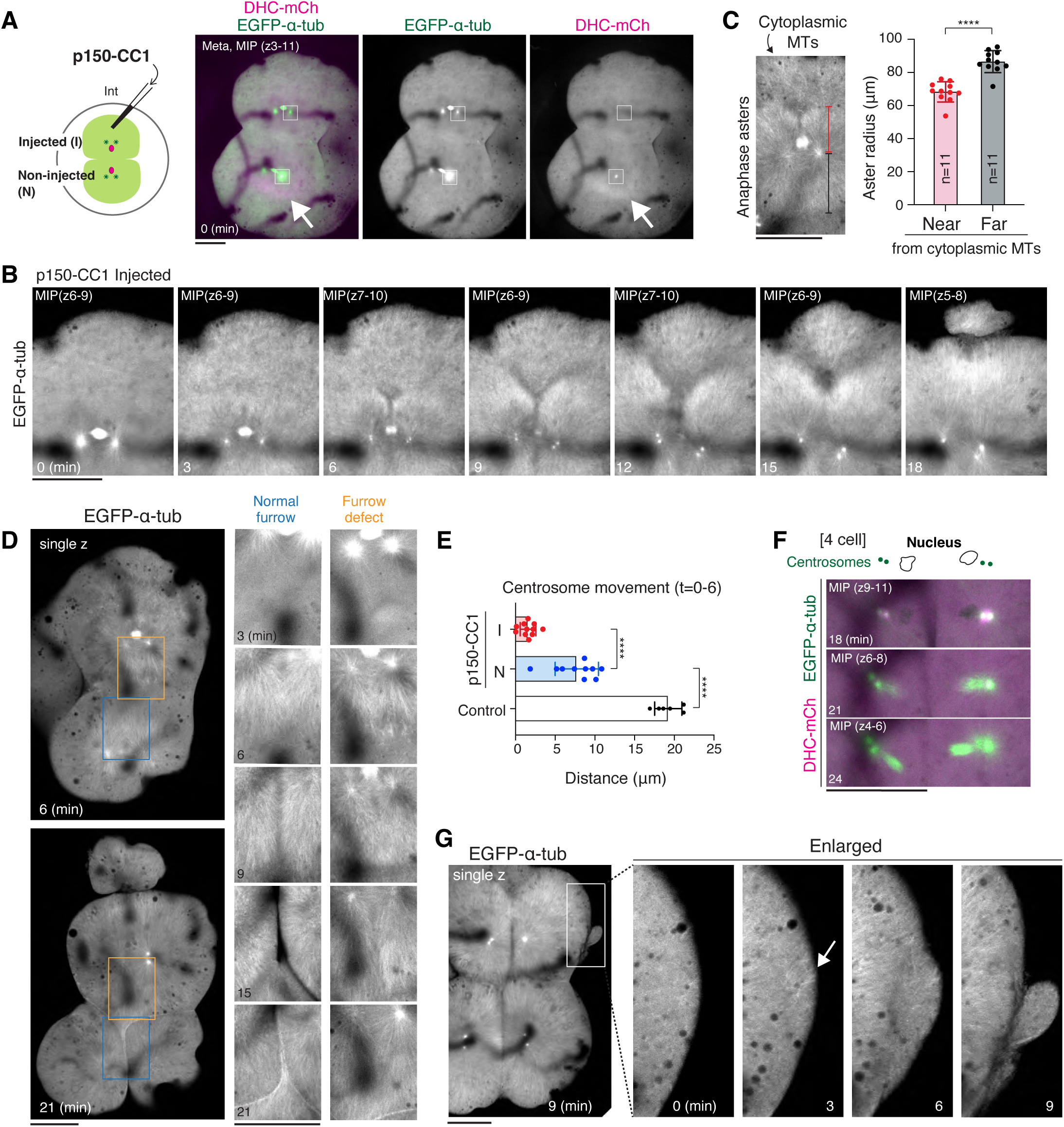
Cytoplasmic MT nucleation, ectopic furrow formation, and cytokinesis defects caused by p150-CC1 injection in a 2-cell blastomere. (A) Live images of MTs (green) and dynein (magenta) in 2-cell blastomeres after p150-CC1 injection. Rectangular regions are enlarged in Fig. 4C. Arrows indicate the dynein halo in the non-injected blastomere. (B) Time-lapse images of p150-CC1-injected 2-cell blastomere showing gross cytoplasmic MT nucleation in anaphase (t=3). MT-less Y-shaped region was created between the cytoplasmic MTs and asters (t=9-12). Eventually, aster growth induced an ectopic furrow (t=18). (C) Left: EGFP-α-tubulin image of p150-CC1-injected blastomere showing that aster radius became shorter near the cytoplasmic MTs. Right: Graphs showing aster radius near or far from cytoplasmic MTs. ****p<0.0001 (D) Left: single-z images of MTs (EGFP-α-tubulin) in another p150-CC1-injected 2-cell embryo showing that cleavage furrow is not formed between asters despite aster growth in p150-CC1-injected blastomere (orange rectangles), whereas furrow is created in the other non-injected-blastomere (blue rectangles). Right: enlarged time-lapse images of MTs in the indicated regions. (E) Graphs showing distances of centrosome movement in p150-CC1 injected blastomeres (red, n=11), non-injected sister-blastomeres (blue, n=10), and control (2-cell blastomeres of non-injected embryos, white, n=6). ****p<0.0001 (F) Live images of MTs (green) and dynein (magenta) at the 4-cell stage of non-injected sister-blastomere showing that abnormal nucleus-centrosome positioning in interphase (t=18) lead to abnormal spindle formation in the subsequent mitosis (t=24). The whole embryo images are shown in Fig. 4B. (G) Single z-section images of MTs after injection of diluted p150-CC1 showing that cytoplasmic MT nucleation near the cell cortex (arrow, t=3) caused a bud-like ectopic furrow (t=9). Scale bars = 100 μm.

**Figure S9.**
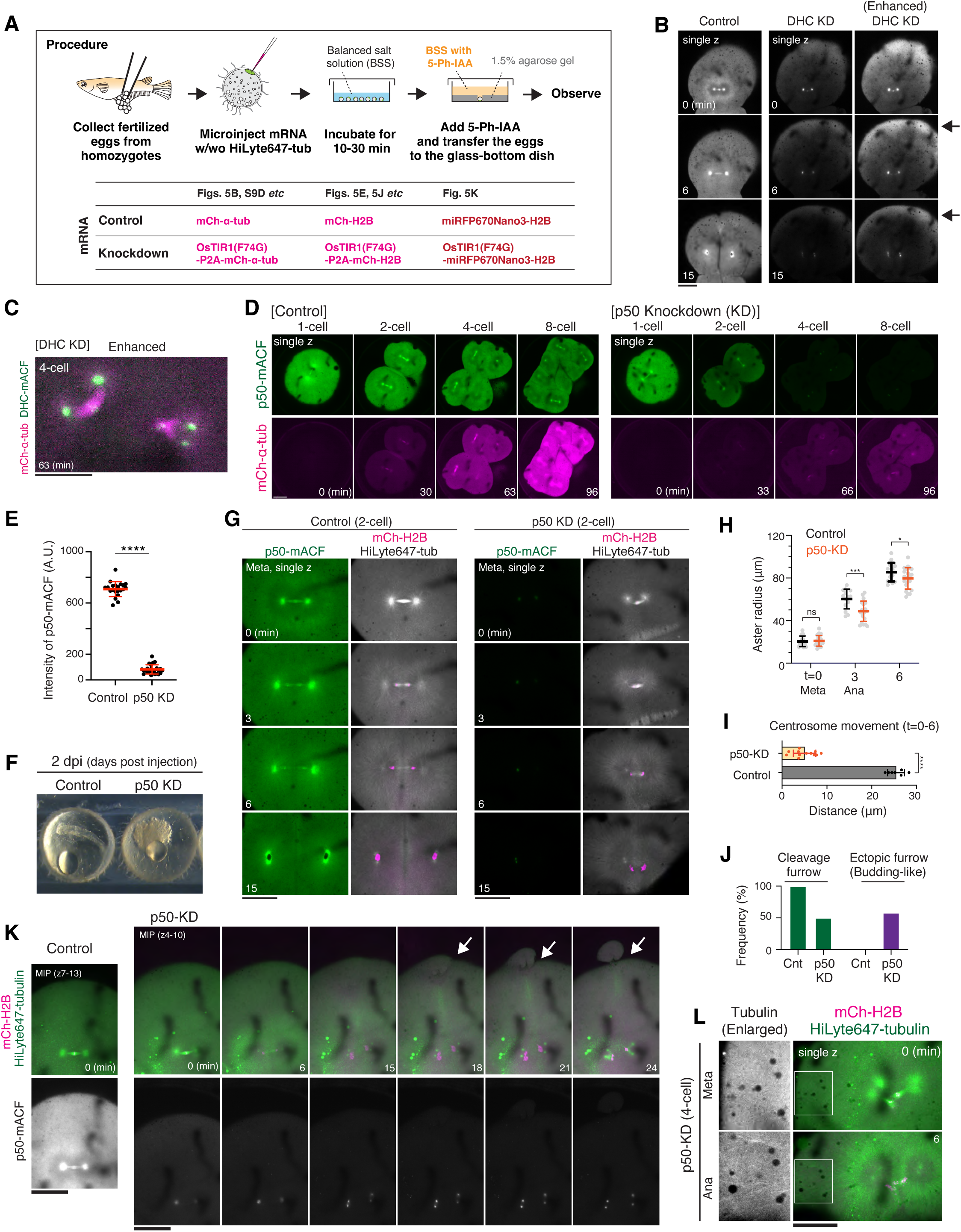
Mitotic phenotypes caused by AID-mediated dynein or dynactin knockdown in medaka early embryos. (A) Procedure of AID-mediated protein knockdown in medaka fertilized eggs. For knockdown, OsTIR-F74G were simultaneously expressed with mCh-α-tubulin or mCh-H2B by connecting them with P2A sequence. Both control and knockdown embryos were incubated with 10 μM 5-PhIAA and observed in the same dish. (B) Live fluorescence images of DHC-mACF in control and DHC-knockdown (KD) 2-cell-stage embryos. DHC-mACF intensities at the cell periphery (arrows) tended to slowly decrease compared to those around the cell center. (C) Live image of DHC-KD embryo showing abnormal spindles at the 4-cell stage. Fluorescent intensities of DHC-mACF (green) and MTs (mCh-α-tubulin, magenta) were enhanced. (D) Representative live-cell images showing the fluorescence of p50-mACF and mCh-α-tubulin in control (left) and OsTIR1(F74G)-expressing (right) embryos. In our system shown in A, mCh-α-tubulin intensities in p50-KD embryos are lower than those in control due to P2A-mediated expression. (E) Quantification of fluorescence intensity of p50-mACF one frame before 4-cell mitosis in control (n=21) and p50-KD (n=24) blastomeres. (F) A phase contrast image showing lethality in a p50-KD embryo (right). (G) Live-cell images of control (left) and p50 KD 2-cell blastomeres showing chromosomes (mCh-H2B, magenta) and MTs (HiLyte647-tubulin, white). mRNAs encoding either miRFP670Nano3-α-tubulin or OsTIR1-F74F-P2A-miRFP670Nano3-α-tubulin were co-injected with HiLyte647-tubulin in control or p50-KD embryos, respectively. Since miRFP670Nano3-α-tubulin did not show any fluorescence in early embryos, MTs were visualized only by HiLyte647-tubulin. Fluorescence intensities of endogenous p50-mACF were adjusted with the same scale between control and p50-KD blastomeres, but those of MTs and chromosomes were differently adjusted since their fluorescence intensities were variable due to injection amount. (H) Graphs showing aster radii in control (n=14) and p50-KD (n=21) blastomeres. *p<0.1, ***p<0.001. (I) Graphs showing distances of centrosome movement in control (n=7) and p50-KD (n=12) blastomeres. ****p<0.0001. (J) Frequencies of a normal cleavage furrow and bud-like ectopic furrows in control (n=7) and p50-KD (n=12) 2-cell stage embryos. (K) Live images showing a normal furrow in control (left) and a bud-like ectopic furrow (arrows) in a p50-KD blastomere (right). (L) Another example of anaphase cytoplasmic MT nucleation caused by p50-KD in 4-cell blastomeres. Scale bars = 100 μm.

**Figure S10.**
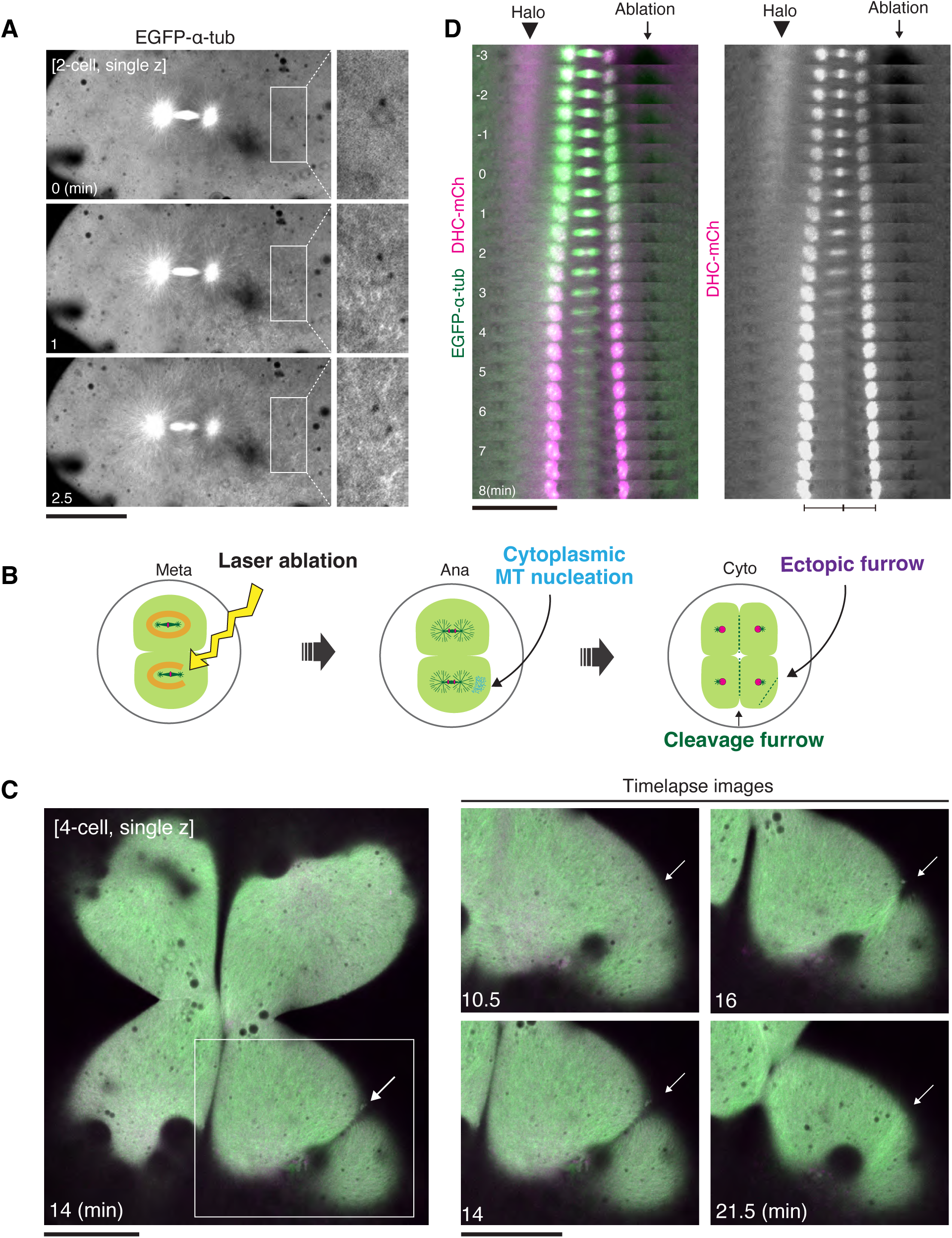
Local disruption of the dynein halo by laser ablation causes asymmetric cytoplasmic MT nucleation. (A) Live images of MTs (EGFP-α-tubulin) showing cytoplasmic MT nucleation in anaphase near the ablated region. (B) Diagrams showing that local disruption of the dynein halo causes cytoplasmic MT nucleation in anaphase, leading to ectopic furrow formation. (C) Left: a live single z-section image of the 4-cell-stage embryo used in Fig. 6B showing an ectopic furrow after laser ablation (arrow, t=14). The region indicated with a white rectangle is enlarged on the right. Right: time-lapse images of MTs (green) and dynein (magenta) showing pseudo cleavage of the induced furrow. (D) Kymographs showing the dynamics of the dynein halo and centrosomes after local halo disruption by laser ablation. The right side of the dynein halo was locally disrupted during metaphase, which attenuated outward centrosome movement after anaphase. Scale bars = 100 μm.

**Figure S11.**
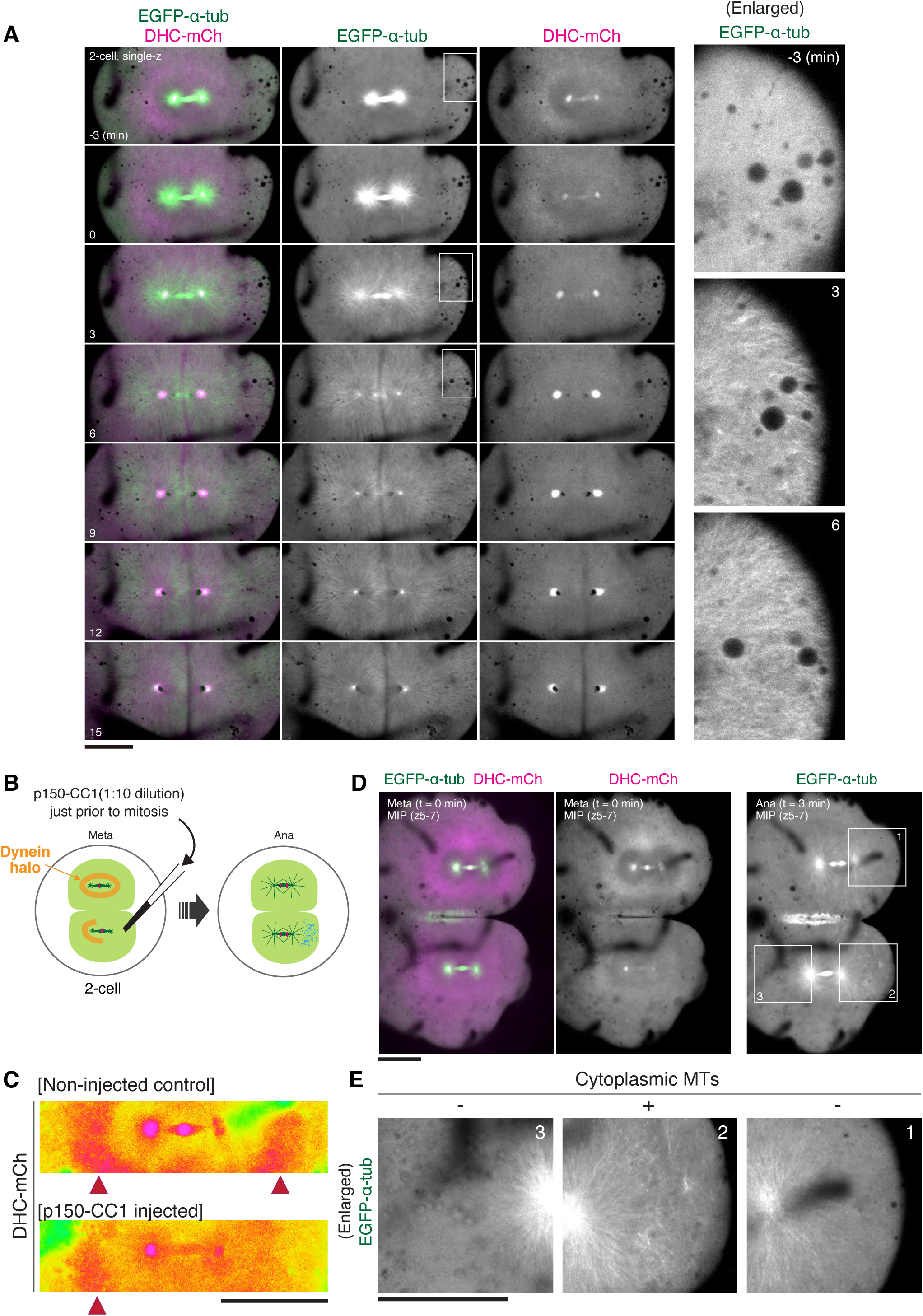
Mitotic phenotypes caused by asymmetric halo disruption resulting from diluted p150-CC1 injection. (A) Time-lapse images of MTs (EGFP-α-tubulin, green) and endogenous dynein (DHC-mCh, magenta) in the 2-cell stage blastomere shown in Fig. 6G. The indicated regions with white rectangles are enlarged on the right. Asymmetric disruption of the dynein halo caused asymmetric cytoplasmic MT nucleation in anaphase (t=3). (B) Schematics showing phenotypes after diluted p150-CC1 injection in one blastomere of a 2-cell embryo. (C) Heatmap images of dynein (DHC-mCh) in another embryo showing an asymmetric dynein halo created by diluted p150-CC1 injection (bottom). The non-injected sister blastomere (control) shows symmetric dynein halo (top). (D) Live images of dynein (DHC-mCh, magenta) and MTs (EGFP-α-tubulin, green) of the 2-cell stage embryo showing asymmetric dynein halo disruption followed by asymmetric cytoplasmic MT nucleation in anaphase. (E) Enlarged images of indicated regions in D showing asymmetric cytoplasmic MT nucleation caused by asymmetric dynein-halo disruption. Scale bars = 100 μm.

## Notes

### Competing Interest Statement

The authors have declared no competing interest.

